# Itk promotes the integration of TCR and CD28 costimulation, through its direct substrates, SLP-76 and Gads

**DOI:** 10.1101/2020.09.11.293316

**Authors:** Enas Hallumi, Rose Shalah, Wan-Lin Lo, Jasmin Corso, Ilana Oz, Dvora Beach, Samuel Wittman, Amy Isenberg, Meirav Sela, Henning Urlaub, Arthur Weiss, Deborah Yablonski

## Abstract

The costimulatory receptor, CD28, synergizes with the T cell antigen receptor (TCR) to promote IL-2 production, cell survival and proliferation. Despite their profound synergy, the obligatory interdependence of the signaling pathways initiated by these two receptors is not well understood. Upon TCR stimulation, Gads, a Grb2-family adaptor, bridges the interaction of two additional adaptors, LAT and SLP-76, to form a TCR-induced effector signaling complex. SLP-76 binds the Tec-family tyrosine kinase, Itk, which phosphorylates SLP-76 at Y173 and PLC-γ1 at Y783. Here we identified Gads Y45 as an additional TCR-inducible, Itk-mediated phosphorylation site. Y45 is found within the N-terminal SH3 domain of Gads, an evolutionarily conserved domain with no known binding partners or signaling function. Gads Y45 phosphorylation depended on the interaction of Gads with SLP-76 and on the preferentially-paired binding of Gads to phospho-LAT. Three Itk-related features, Gads Y45, SLP-76 Y173, and a proline-rich Itk SH3-binding motif on SLP-76, were selectively required for activation of the CD28 RE/AP transcriptional element from the IL-2 promoter, but were not required to activate NFAT. This study illuminates a new regulatory module, in which Itk-targeted phosphorylation sites on two adaptor proteins, SLP-76 and Gads, control the transcriptional response to TCR/CD28 costimulation, thus enforcing the obligatory interdependence of the TCR and CD28 signaling pathways.

## 1. Introduction

The TCR signaling pathway [recently reviewed in 1, 2, 3] is initiated by a hierarchical tyrosine kinase cascade, leading to the formation of a large effector signaling complex that is nucleated by three interacting adaptor proteins, LAT, Gads and SLP-76 (Figure 1). Within this LAT-nucleated complex, adaptor-associated enzymes are recruited to become activated and trigger downstream responses. For example, phospholipase-Cγ1 (PLC-γ1) is a key signaling enzyme that is phosphorylated and activated within the LAT-nucleated complex and produces second messengers, IP3 and DAG, which respectively trigger calcium flux and Ras-pathway activation. Further downstream, these signaling events result in the activation of well-characterized transcription factors, including NFAT and AP-1.

**Figure 1.**
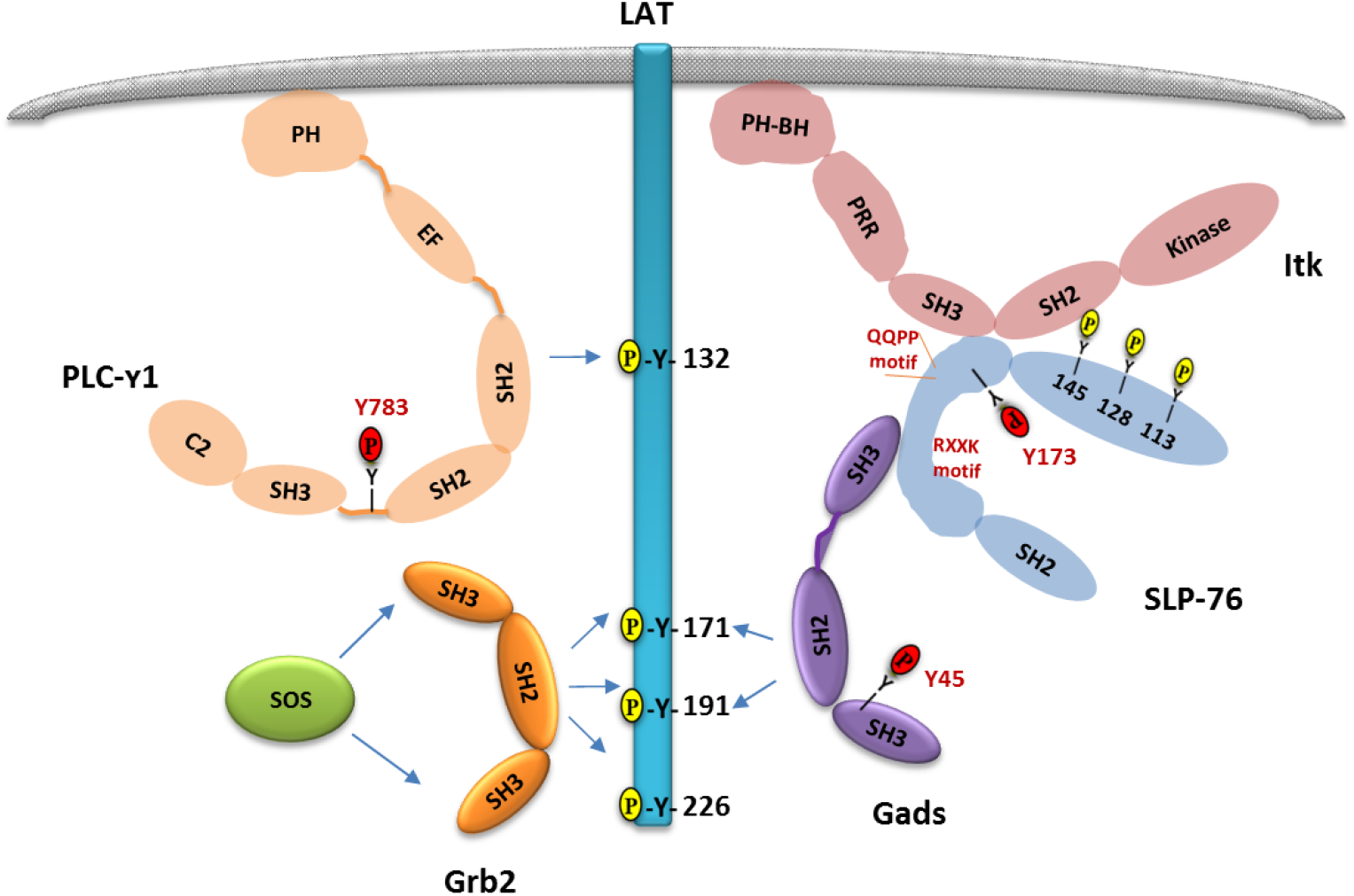
A web of interactions connects Itk to its substrates. Upon TCR stimulation, ZAP-70 phosphorylates conserved tyrosine residues on LAT and SLP-76 (shown in yellow). Itk binds to SLP-76 via a multivalent interaction, in which its SH2 may bind to SLP-76 p-Y145, and its SH3 domain may bind to the SLP-76 QQPP motif. Gads bridges the recruitment of SLP-76 to LAT, via its C-SH3 domain, which binds with high affinity to a SLP-76 RxxK motif, and its central SH2 domain, which binds in a cooperatively paired manner to LAT p-Y171 and p-Y191. Grb2 bridges the recruitment of SOS to LAT p-Y171, p-Y191 and/or p-Y226. PLC-γ1 is recruited via its N-SH2 to LAT p-Y132, and is thereby brought into the vicinity of SLP-76-bound Itk. Itk-targeted sites, including Gads Y45, identified in this study, are shown in red.

In addition to the above events, full T cell responsiveness depends on costimulatory signals that are triggered upon binding of B7-family ligands to the canonical costimulatory receptor, CD28. The TCR and CD28 signaling pathways act synergistically to induce the activation of NFκB-family members, which are required for the transcriptional activation of IL-2 and Bcl-xL, key response markers that promote T cell proliferation and survival [4, 5].

The TCR and CD28 signaling pathways exhibit remarkable interdependence, which can be most easily demonstrated by measuring the activity of RE/AP, a composite transcriptional element that forms an essential part of the IL-2 promoter [6, 7]. Composed of adjacent AP-1 and NFκB sites that bind to c-Jun and to c-Rel, the RE/AP element recapitulates the costimulation-dependence of IL-2 transcription, as stimulation with either TCR or CD28 alone is insufficient to activate RE/AP, but co-stimulation through both receptors produces profound RE/AP activation [7, 8].

The interdependence of TCR and CD28 responsiveness has important biological outcomes. CD28 costimulatory ligands serve as “danger signals” to indicate the presence of microorganisms that necessitate an adaptive immune response [9]. Yet, to avoid the induction of potentially dangerous bystander activity, CD28-dependent T cell activation must necessarily be restricted to those cells bearing an antigen-specific clonotypic TCR. This goal is achieved by profound interdependence, such that CD28 activity is necessary but insufficient for the activation of naive T cells. However, the mechanistic basis for TCR/CD28 interdependence is not well understood.

The basic outlines of the TCR signaling pathway are well-defined [reviewed in 1, 10]. Upon TCR stimulation, a tyrosine kinase cascade is initiated by Lck, a Src-family tyrosine kinase that phosphorylates characteristic ITAM motifs within the CD3 and ζ accessory chains of the TCR. Doubly-phosphorylated ITAM motifs trigger the recruitment and activation of a Syk-family tyrosine kinase, ZAP-70 [11]. Activated ZAP-70 phosphorylates LAT, a membrane-bound adaptor, on at least four essential sites [12–15], and phosphorylates SLP-76, a cytoplasmic adaptor, at three N-terminal phosphorylation sites [Figure 1 and 16, 17]. The ZAP-70-phosphorylated sites recruit additional signaling molecules via SH2-mediated interactions. Phospho-LAT binds directly to PLC-γ1, Grb2 and Gads, whereas the three tyrosine phosphorylation sites on SLP-76 bind to Nck, Vav and Itk [reviewed in 2, 18].

Itk, a Tec-family tyrosine kinase, is the third member of the TCR-induced tyrosine kinase cascade, as it is activated downstream of both Lck and ZAP-70. Lck phosphorylates Itk at Y511 [19], whereas ZAP-70 phosphorylates SLP-76 at three N-terminal tyrosines that are required for Itk activation [16, 17, 20, 21]. Upon TCR stimulation, Itk associates with SLP-76, and this association is required to maintain the active conformation of the kinase [20]. SLP-76-bound Itk phosphorylates SLP-76 at Y173 [21]. In turn, SLP-76 p-Y173 promotes the subsequent Itk-mediated phosphorylation of PLC-γ1 at Y783 [21, 22], a site that is required for PLC-γ1 activation [23].

Structural studies suggest that an inactive conformation of Itk is stabilized by intramolecular interactions of its SH2 and SH3 domains, as well as by an inhibitory interaction between the N-terminal PH domain and the kinase domain [24–26]. Upon TCR or CD28 stimulation, PIP3 in the plasma membrane is increased. Elevated PIP3 binds to the Itk PH domain, relieving the inhibitory influence of the PH domain on the kinase domain [26]; moreover, bivalent binding of the SH2 and SH3 domains of Itk to SLP-76 is thought to stabilize the active confirmation of Itk [Figure 1, right and 27, 28, 29].

The SH2 domain of Itk is thought to bind to SLP-76 p-Y145 [25]; however, competitive binding studies suggested that it may also bind to SLP-76 p-Y113 [28], which is equivalent to murine p-Y112. A SLP-76 Y145F mutation closely phenocopied Itk-deficient mice, but was insufficient to eliminate the TCR-induced binding of Itk to SLP-76 [30]. Unexpectedly, phosphorylation PLC-γ1 Y783 was markedly reduced by either the SLP-76 Y145F mutation, or by the double SLP-76 Y112,128F mutation [21]. Thus, although Itk is commonly thought to bind to SLP-76 Y145 [25, 31], evidence suggests that at least two N-terminal tyrosines of SLP-76 are required, whether directly or indirectly, for optimal activation of Itk.

The SH3 domain of Itk can bind to a conserved proline rich motif, QQPPVPPQRP, corresponding to SLP-76 residues 184-195 [28, 29], which, for convenience, we shall refer to as the QQPP motif. This motif was also reported to bind, albeit weakly, to the SH3 domains of Lck and PLC-γ1 [32–35]. Precise removal of the QQPP motif in a transgenic mouse model reduced TCR-stimulated PLC-γ1 phosphorylation and calcium flux, as would be expected if Itk activity were disrupted [34, 36]. Moreover, a cell permeable peptide based on the QQPP motif inhibited the TCR-inducible interaction of Itk with SLP-76, phosphorylation of Itk at Y511, recruitment of Itk to the immune synapse, and consequently inhibited the production of Th2 cytokines [29], all consistent with a defect in Itk-mediated signaling [36, 37]. Nevertheless, deletion of a 36 amino acid region encompassing the QQPP motif only modestly reduced PLC-γ1 phosphorylation and calcium flux in reconstituted J14 cells [32], suggesting that the role of the QQPP motif in regulating Itk activity may be context-dependent.

Itk-mediated phosphorylation of PLC-γ1 is facilitated by Gads [38, 39], a Grb2-family adaptor that bridges the TCR-inducible recruitment of SLP-76 to phospho-LAT [Figure 1A, center, reviewed in 3]. Composed of a central SH2 domain, flanked by two SH3 domains and a unique linker region, Gads binds constitutively to SLP-76, via a high affinity interaction of its C-terminal SH3 domain with a conserved RxxK motif on SLP-76 [33, 40, 41]. The SH2 domain of Gads is capable of spontaneous dimerization, and mediates the cooperatively-paired binding of Gads to LAT p-Y171 and p-Y191, thereby recruiting SLP-76 to phospho-LAT [42]. Curiously, the bridging activity of Gads does not require its N-terminal SH3 domain, an evolutionarily conserved domain that has no known ligand or signaling function [3]. Gads bridging activity supports PLC-γ1 phosphorylation, by bringing SLP-76-bound Itk (Figure 1, right) in proximity with its substrate, LAT-bound PLC-γ1 (Figure 1, left).

Once formed, the LAT-nucleated signaling complex may be regulated by post-translational events occurring within the complex. HPK1, a SLP-76-associated Ser/Thr kinase, can negatively regulate TCR signaling by phosphorylating SLP-76 at S376 [43, 44], and by phosphorylating Gads at T262 [38, 45]. Conversely, SLP-76-associated Itk promotes TCR responsiveness by phosphorylating SLP-76 at Y173 [21]. Itk activity is highly dependent on docking interactions that target its catalytic activity to potential substrates [22, 46, 47]. Since Itk is inducibly docked onto SLP-76, which binds with high affinity to Gads, this paradigm suggested to us that additional Itk-mediated phosphorylation sites on SLP-76 or Gads might play an important role in regulating their signaling function.

To explore this hypothesis, we performed a phospho-mass spectrometry analysis of SLP-76 and Gads, which were isolated from TCR-stimulated cells. Here we identify Gads Y45 as a TCR-inducible substrate of Itk, which is phosphorylated within the SLP-76-Gads-LAT signaling complex. Y45 is found within the N-terminal SH3 domain of Gads, and may provide a first clue to the biological function of this conserved domain. Unexpectedly, we found that TCR/CD28-induced activation of the RE/AP transcriptional element depended on Gads Y45 and SLP-76 Y173, two Itk-targeted sites, and also depended on the QQPP motif, an Itk-binding site within SLP-76. Gads Y45 phosphorylation was strictly dependent on the TCR-induced cooperative binding of Gads to LAT, and thereby may enforce the dependence of CD28 responsiveness on TCR activation.

### 2. Materials and Methods

#### 2.1. Recombinant Gads proteins

Recombinant, maltose-binding protein (MBP)-Gads fusion proteins were expressed and purified as previously described [42].

#### 2.2. Antibodies

To prepare mouse anti-Gads, Gads deficient mice on the Balb/C background were immunized with recombinant MBP-Gads-ΔN-SH3 in Freund’s adjuvant. Polyclonal, affinity-purified rabbit anti-phospho-Gads p-Y45 was prepared for us by GenScript, by immunizing rabbits with a phospho-peptide GSQEG{p-TYR}VPKNFIDIC, corresponding to amino acids 40–54 of human Gads, conjugated to KLH, followed by two steps of affinity chromatography, to remove antibodies that recognize the non-phosphorylated peptide and enrich for those that recognize the phosphorylated peptide. To decrease non-specific background without blocking the sequence-specific recognition of p-Y45, we supplemented the diluted, purified antiserum with 3 μg/ml phosphotyrosine-conjugated BSA (Sigma, P3967). Rabbit polyclonal anti-human SLP-76 [32] and rabbit anti phospho SLP-76 p-Tyr173 [21] were previously described. Polyclonal anti-Itk [BL12, 48] was provided by Michael G Tomlinson and Joseph Bolen and was used for immunoprecipitation. The monoclonal antibody C305 [49] was used to stimulate Jurkat-derived cell lines through the TCR. Purified anti-human CD28 clone CD28.2, anti-human CD3-APC, anti-human CD28-PE and anti-human CD69 PerCP/Cy5.5 (clone FN50) were from Biolegend. Rabbit anti-PLC-γ1 (sc-81) was from Santa Cruz Biotechnology. Rabbit anti-phospho PLC-γ p-Tyr783 (AT-7142) was from MBL International. Rabbit polyclonal anti-phospho LAT p-Tyr191 (#3584) was from Cell Signaling Technology. Anti p-Tyr (4G10) and anti-Itk (06-546, used for western blotting) were from Merck Millipore. Anti-phospho Itk p-Tyr511 (clone M4G3LN) was from ThermoFisher. Anti-phosho LAT p-Tyr171 (clone I58-1169) was from BD Biosciences. Anti-phosho Jnk pT183+pT221-PE (ab208843) was from Abcam. Anti-Human IL-2-PE (MQ1-17H12) was from eBioscience.

#### 2.3. Cell lines and retroviral reconstitution

Cell lines used in this work are summarized in Table 1. Cells were grown in RPMI, supplemented with penicillin, streptomycin and glutamine (PSG) and 5% fetal calf serum (FCS) in a humidified incubator with 5% CO2. Cells were retrovirally reconstituted with wild-type or mutant, N-terminally twin-strep tagged [50] or FLAG-tagged, human Gads or SLP-76, using the pMIGR retroviral vector, which bears an internal ribosome entry site (IRES)-GFP cassette to mark infected cells [51]. Approximately two weeks later cells were sorted by FACS for comparable GFP expression, and comparable TCR and CD28 expression were verified by FACS. Where indicated, we used a modified, IRES-less version of pMIGR [42] in which Gads was fused C-terminally to a non-dimerizing form of GFP [GFP A206K, 52].

**Table 1.**
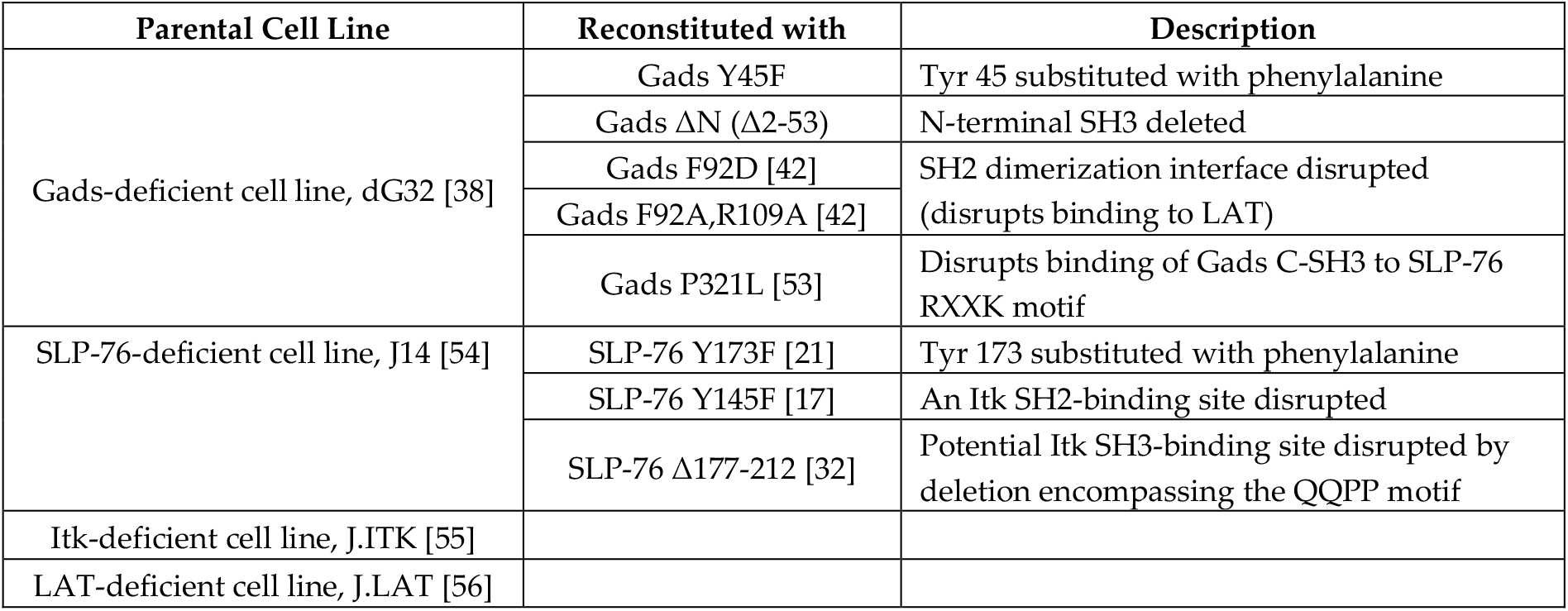
A summary of the cell lines and point mutations employed in this study.

| Parental Cell Line | Reconstituted with | Description |
| --- | --- | --- |
| Gads-deficient cell line, dG32 [38] | Gads Y45F | Tyr 45 substituted with phenylalanine |
| | Gads ΔN (Δ2-53) | N-terminal SH3 deleted |
|  | Gads F92D [42] | SH2 dimerization interface disrupted (disrupts binding to LAT) |
|  | Gads F92A,R109A [42] |  |
|  | Gads P321L [53] | Disrupts binding of Gads C-SH3 to SLP-76 RXXK motif |
| SLP-76-deficient cell line, J14 [54] | SLP-76 Y173F [21] | Tyr 173 substituted with phenylalanine |
|  | SLP-76 Y145F [17] | An Itk SH2-binding site disrupted |
| | SLP-76 Δ177-212 [32] | Potential Itk SH3-binding site disrupted by deletion encompassing the QQPP motif |
| Itk-deficient cell line, J.ITK [55] |  |  |
| LAT-deficient cell line, J.LAT [56] |  |  |

#### 2.4. Cell stimulation and Purification of Gads or SLP-76 complexes

Jurkat-derived T cell lines were washed and resuspended in stimulation medium (RPMI + 100 μg/ml glutamine), preheated to 37°C for 10 min, and then stimulated for the indicated time at 37°C with anti-TCR antibody (C305), with or without 1.5-2 μg/ml anti CD28 as indicated. Cells were lysed at 10^8^ cells/ml in ice-cold lysis buffer (20 mM Hepes pH 7.3, 1% Triton X-100, 0.1% *n*-dodecyl-β-D-maltoside (Calbiochem) 150 mM NaCl, 10% glycerol, 10 mM NaF, 1 mM Na3VO4, 10 μg/ml aprotinin, 2 mM EGTA, 10 μg/ml leupeptin, 2 mM phenylmethanesulfonyl fluoride, 1 μg/ml pepstatin, and 1 mM dithiothreitol). Naïve CD4 T cells were sorted by BD FACSAria II. Thymocytes or purified naïve CD4 T cells from 4-6 week old C57BL/6 mice were stimulated with anti-CD3 (145-2C11) and anti-CD28 (clone 37.51) at the indicated concentration, followed by cross-linking with anti-hamster IgG. Cells were lysed by directly adding 10% NP-40 lysis buffer to a final concentration of 1% NP40 (containing the inhibitors: 2 mM NaVO4, 10 mM NaF, 5 mM EDTA, 2 mM PMSF, 10 μg/ml Aprotinin, 1 μg/ml Pepstatin and 1 μg/ml Leupeptin). Cell lysates were placed on ice and centrifuged at 16,000 × g for 10 min at 4°C to pellet cell debris, and Gads or SLP-76 complexes were affinity purified by tumbling lysates end-over-end for 1-3 hours at 4°C with Strep-Tactin Superflow high capacity beads (IBA), FLAG M2 magnetic beads (Sigma), or with anti-SLP-76, prebound to protein A sepharose fast flow (GE Healthcare). After three rapid washes with cold lysis buffer, the isolated complexes were analyzed by mass spectrometry (see below) or by Western blotting. Western blots were developed with SuperSignal™ West Pico PLUS Chemiluminescent Substrate and digital images of the membrane were produced by the ImageQuant LAS 4000 or the Fusion Fx7 camera system, followed by quantification with Total-Lab Quant software.

#### 2.5. A SILAC-based kinetic analysis of TCR-induced SLP-76 and Gads phosphorylation sites

TCR-inducible phosphorylation sites within the SLP-76-nucleated complex were quantitatively identified by using a Stable Isotope Labeling with Amino Acids in Cell Culture (SILAC)-based approach, exactly as we previously described [38]. In brief, J14 cells, stably reconstituted with twin-strep-tagged SLP-76, were metabolically labeled with the heavy amino acids L-tyrosine ^13^C9^15^N (+10), L-Lysine ^13^C6^15^N2 (+8) and L-Arginine ^13^C6^15^N4 (+10), or with the corresponding light amino acids. Prior to lysis, SILAC-labeled cells were stimulated with anti-TCR (heavy label) or mock-stimulated (Iight label). Heavy and light lysates derived from 120 million cells were mixed at a 1:1 ratio, followed by affinity purification of twin-strep-tagged SLP-76 and its associated proteins with Strep-Tactin beads. Purified proteins were split into three samples that were analyzed in parallel. Following SDS-PAGE, Coomassie-stained protein bands corresponding to Gads and SLP-76 were cut from the gel, followed by in gel digestion with trypsin, chymotrypsin or AspN. Phospho-peptides were enriched with TiO2 and were analyzed on an LTQ Orbitrap Velos (Thermo Fischer) mass spectrometer coupled to a nanoflow liquid chromatography system (Agilent 1100 series, Agilent), as described [57]. Resulting raw files were processed with MaxQuant (v1.3.0.5) against a UniProtKB/Swiss-Prot human database. Data presented are the median values from four biological repeats.

#### 2.6. Kinase assays

Polyclonal anti-Itk (BL12) was used to immunoprecipitate Itk from the lysates of 4 × 10^6^ TCR-stimulated dG32 cells. IP beads were washed twice with lysis buffer and once with kinase reaction buffer (25 mM HEPES pH 7.3, 7.5 mM MgCl_2_, and 1 mM Na_3_VO_4_), resuspended in 30 μl of kinase reaction buffer containing 1 μM of recombinant MBP-Gads protein, and preheated for 2 min at 30 °C. Kinase activity was initiated by the addition of ATP to 1 μM, and was terminated after 30 min at 30 °C with end-over-end mixing, by the addition of EDTA to 12.5 mM.

#### 2.7. FACS-based functional assays

To decrease experimental variation, cell lines were barcoded by differential labeling with fourfold dilutions of CellTrace Far Red or CellTrace Violet (ThermoFisher), and mixed together prior to stimulation as described [42]. For calcium assays, mixed, CellTrace Far Red-barcoded cells were loaded with the fluorescent calcium indicator dye, Indo1-AM (eBioscience), and then washed twice and resuspended in calcium buffer, consisting of 25 mM Hepes (pH 7.4), 1 mM CaCl_2_, 125 mM NaCl, 5 mM KCl, 1 mM Na_2_HPO_4_, 0.5mM MgCl_2_, 1 g/l glucose, 2 mM L-glutamine and 1 mg/ml high-purity bovine serum albumin (Sigma A4378). Intracellular calcium was measured by FACS at 37°C, with C305 or C305+CD28.2 stimulant added at the 60s time point [38]. CellTrace Violet-barcoded cells were stimulated as indicated in the figure legends, and stained with anti-CD69, or were fixed, permeabilized and stained with anti-pJNK, or with anti-IL-2-PE (eBioscience). Results were analyzed using FlowJo, while gating on a defined GFP window within each barcoded population.

#### 2.8. Luciferase Assays

The firefly luciferase reporter plasmids pdelta-ODLO-3XNFAT and pdelta-ODLO-4XRE/AP [7] were provided by Virginia Shapiro (Mayo Clinic). 5xκB-luciferase was from Stratagene. pRL-null, which drives constitutive expression of renilla luciferase (Promega), was used for normalization. 20X10^6^ cells were transfected by electroporation with 20 μg of firefly luciferase reporter plasmid and 3-5 μg of pRL-null, using the Gene Pulser (Bio-Rad Laboratories), at a setting of 234 V and 1000 microfarads. 16-20 hr after transfection 2X10^5^ cells per well were stimulated for 6 hr at 37°C in a 96 well plate format with plate-bound anti-TCR (C305) and/or soluble anti-CD28 (CD28.2 1.5 μg/ml) or were mock-stimulated, and activity was measured with the Dual Luciferase Kit (Promega). To correct for variations in transfection efficiency, firefly luciferase activity for each well was normalized the renilla luciferase activity measured in the same well.

### 3. Results

#### 3.1. TCR-inducible phosphorylation of Gads at Y45

We previously described a SILAC-based approach to identify TCR-inducible phosphorylation sites within the SLP-76-nucleated complex [38]. Here, we used this approach to survey the TCR-inducible phosphorylation sites on SLP-76 and Gads. Of the sites we identified, peptides harboring SLP-76 p-Y173 and Gads p-Y45 exhibited the highest fold increase in intensity upon TCR stimulation (Figure S1A and S1B). We previously characterized SLP-76 p-Y173, a TCR-inducible site that is phosphorylated by Itk and facilitates the subsequent Itk-mediated phosphorylation of PLC-γ1 [21]; in contrast, the regulation and function of Gads p-Y45 are completely unknown.

We were intrigued by the high fold-induction of Gads p-Y45 upon TCR stimulation (Figure 2A) and by the evolutionary conservation of the sequence motif surrounding this site (Figure 2B). Gads Y45 is found within the N-terminal SH3 domain of Gads, which, like the C-terminal SH3 domain, is highly conserved (Figure S2). Whereas the C-SH3 binds with high affinity to SLP-76; the N-SH3 has no known function [3]. Phosphorylation of Gads Y45 was previously observed in high-throughput phospho-MS studies [58–61]; yet its functional significance was not previously explored.

**Figure 2.**
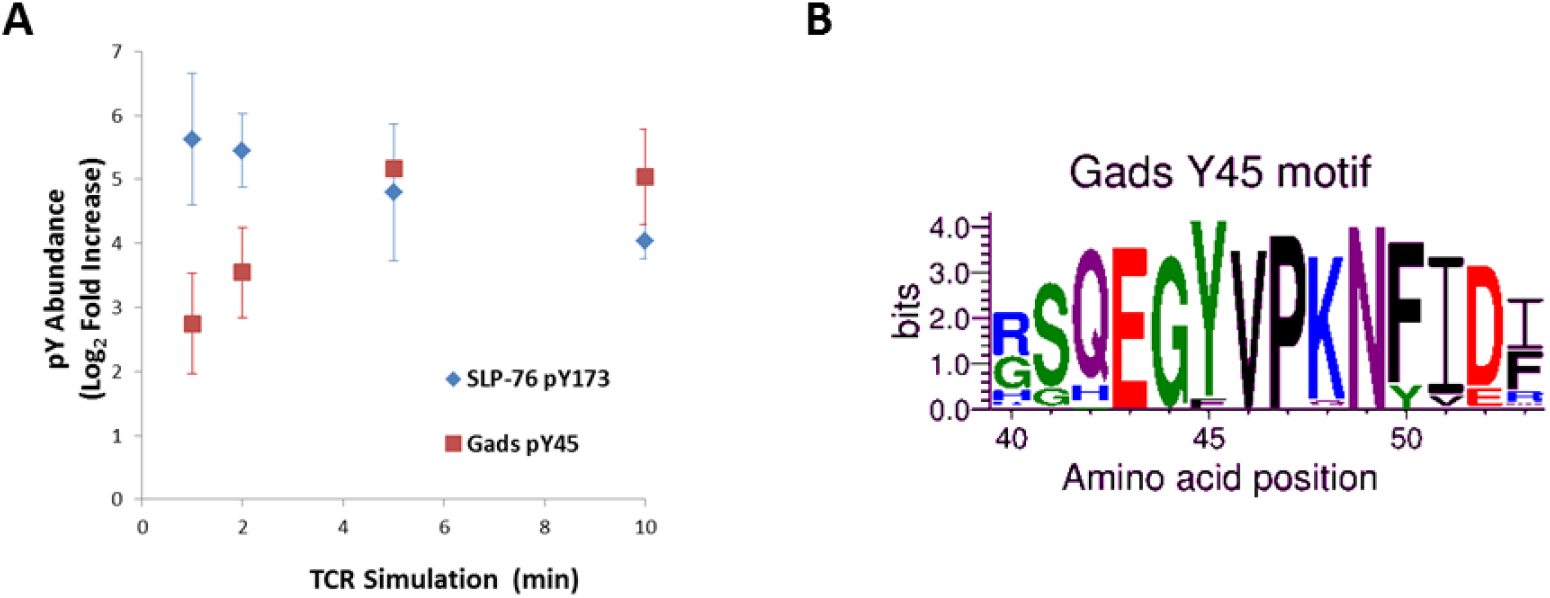
A conserved Gads tyrosine phosphorylation site, identified by MS. (**A**) **Kinetics of the TCR-induced phosphorylation of Gads Y45 and SLP-76 Y173.** A SILAC approach was employed to measure the TCR-induced fold change in SLP-76 and Gads phosphorylation sites. Shown is the median Log2-fold change for the two most highly-induced sites: Gads p-Y45 and SLP-76 p-Y173. Data are from four independent biological replicates; error bars indicate the SD. (**B**) **Evolutionary conservation of the sequence motif surrounding Gads Y45.** NCBI Protein Blast was used to identify and select 66 vertebrate Gads orthologs from the mammalian, avian, cartilaginous and bony fish, amphibian, and reptilian classes, including representatives of 55 different taxonomical orders [3]. Sequences were aligned with Clustal O [62], and WebLogo [63] was used to depict the conservation of Gads residues 40-53.

To fill the gaps in our fundamental knowledge regarding the potential roles of Gads Y45 in TCR signaling, we took advantage of a Gads-deficient T cell line, dG32 [38], which we stably reconstituted with N-terminally twin-strep-tagged Gads, either wild-type (WT) or bearing a phenylalanine substitution at Y45 (Y45F). We also generated an affinity-purified, phospho-specific, polyclonal antibody, to enable us to specifically detect the phosphorylation of Gads Y45.

For routine detection of Gads Y45 phosphorylation, cells were costimulated with anti-TCR and anti-CD28, and strep-tactin-purified Gads complexes, which include Gads-associated SLP-76, LAT and PLC-γ1, were probed by immuno-blotting. TCR/CD28-inducible phosphorylation of Gads was clearly detectable using either the p-Y45 phospho-specific reagent or a global p-Tyr antibody (clone 4G10), and this band was substantially reduced by the Y45F mutation (Figure 3A, top three panels). This result validates our phospho-specific reagent and suggests that Y45 is the major Gads tyrosine phosphorylation site that can be detected by the anti-pTyr antibody, 4G10.

**Figure 3.**
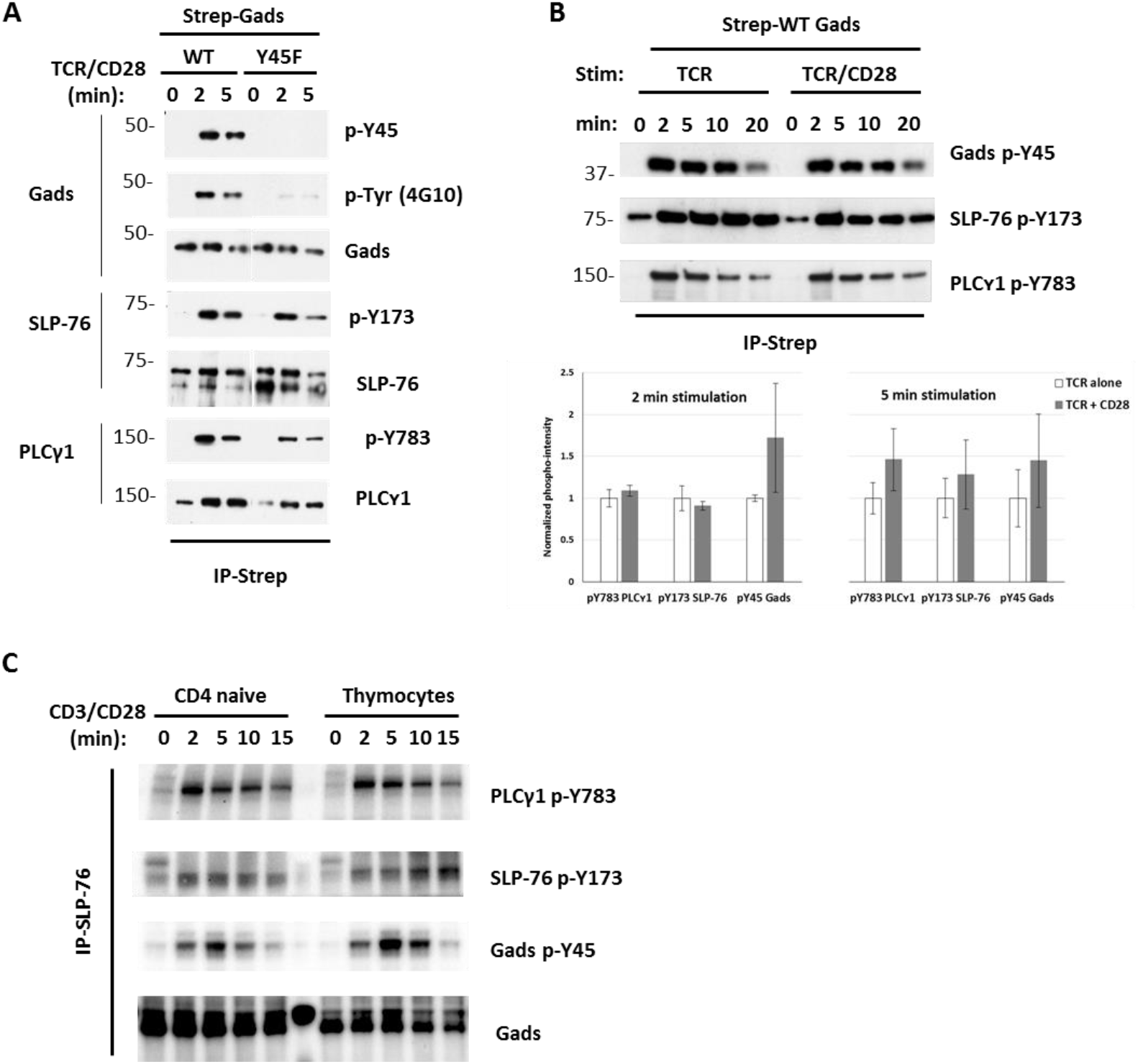
A phospho-specific reagent reveals TCR-induced Gads Y45 phosphorylation in mouse and human T cells. (**A**) **Validation of a p-Y45 phospho-specific reagent.** dG32 cells, stably reconstituted with twin strep-tagged Gads, either WT or Y45F, were stimulated with anti-TCR (C305, 1:1000 dilution) and anti-CD28 (CD28.2 1.5 μg/ml), and then lysed. Gads complexes were isolated with strep-tactin beads (IP-Strep) and probed with the indicated antibodies. Results are representative of at least 5 repeats. (**B**) **Gads Y45 phosphorylation is independent of CD28 co-stimulation.** dG32 cells reconstituted with WT Gads were stimulated with anti-TCR (C305, 1:1000 dilution), in the presence or absence of anti-CD28 (CD28.2 1.5 μg/ml), and lysed. Top: streptactin complexes were probed as in A. Bottom: cells were stimulated for 2 or 5 min in quadruplicate. Streptactin complexes were probed for phospho- and total Gads, whereas lysates were probed for phospho-and total SLP-76 and PLC-γ1, followed by quantification. Phospho-intensity was calculated as the ratio of phospho-to total protein in each band. Results are presented as the fractional phospho-intensity, relative to the average intensity observed in TCR stimulated WT cells from the same stimulation time. (**C**) **Inducible phosphorylation of Gads Y45 in primary mouse T cells.** Prior to lysis, thymocytes or naive CD4 peripheral T cells isolated from C57BL/6 mice were stimulated with 1 μg/ml anti-CD3 and 2 μg/ml anti-CD28, followed by anti-Armenian Hamster IgG crosslinking to induce stimulation for the indicated time at 37°C. SLP-76 complexes were isolated by immunoprecipitation and were probed with the indicated antibodies.

TCR-inducible phosphorylation of Gads Y45 was rapid and sustained, peaking at 2-5 minutes and was still detectable 20 min after stimulation (Figure 3B, top). Phosphorylation of SLP-76 Y173 was similarly rapid and sustained, whereas the phosphorylation of Gads-associated PLCγ1 appeared to be more transient, perhaps reflecting the previously-reported dissociation of phospho-PLCγ1 from LAT [64]. In some experiments, TCR-induced Gads Y45 phosphorylation was moderately augmented by CD28 costimulation, but this difference was not statistically significant (Figure 3B, bottom).

Inducible phosphorylation of Gads Y45 was also observed in primary mouse T cells across their development stages, including thymocytes and naive T cells (Figure 3C). Together, these results demonstrate TCR-inducible phosphorylation of Gads Y45 in both a human T cell line and in primary murine T cells, suggesting a potentially important function, which we set out to investigate.

#### 3.2. Gads Y45 phosphorylation is mediated by Itk

TCR-proximal signaling is mediated by a cascade of three tyrosine kinases, Lck, ZAP-70 and Itk, acting in a strictly hierarchical order. Within this cascade, the activity of each kinase is limited by requirements for characteristic substrate motifs [65] and kinase docking sites [46, 47, 56]. To assess which kinase might be responsible for phosphorylating Gads at Y45, we compared the motif surrounding Gads Y45 (Figure 2B) to the known substrate preferences of Lck, ZAP-70, and Itk. Lck has a strong preference for a bulky hydrophobic residue at the Y-1 position, and does not tolerate lysine at the Y+3 position [65]. Gads Y45 violates both of these rules, suggesting that it is not a substrate of Lck. ZAP-70-targeted sites are characterized by multiple negative residues surrounding the phosphorylated tyrosine; indeed, ZAP-70 is deterred from phosphorylating substrate motifs containing a positive charge anywhere within the surrounding motif [65]. The presence of glycine at the Y-1 position also slows substrate phosphorylation by ZAP-70 [55]. Gads Y45 is preceded by glycine at the Y-1 position, followed by a lysine at the Y+3 position, and has only one negatively charged residue in the surrounding motif, strongly suggesting that it is not a substrate of ZAP-70. To the best of our knowledge, the substrate motifs favored by Itk have not been rigorously defined; however, they are clearly differentiated from ZAP-70 substrates. For example, ZAP-70 efficiently phosphorylates the three N-terminal tyrosines of SLP-76 but does not phosphorylate SLP-76 at Y173, whereas the reverse is true for Itk [21]. Since the conserved motif surrounding Gads Y45 bears some resemblance to known Itk-targeted sites [Fig S3A and 20, 21, 66–69], we decided to test the ability of Itk to phosphorylate Gads Y45, both in an *in vitro* kinase assay, and in an intact cellular environment.

For our *in vitro* experiments, Itk immune complexes were isolated from TCR-stimulated dG32 cells and were incubated in the presence or absence of ATP with recombinant MBP-Gads substrates, either full length (WT), lacking the N-terminal SH3 (ΔN, lacking residues 2-53), or with Phe substituted for Tyr45 (Y45F). All three substrates were phosphorylated by Itk in this assay system, as detected by immunoblotting with global anti-p-Tyr (Figure 4A, 2nd panel). Specific phosphorylation of Gads at Y45 was detected by the p-Y45 antibody, which, as expected, did not recognize the two substrates lacking this site (Figure 4A, top panel). Phosphorylation at all sites was abrogated upon addition of the Itk-specific inhibitor BMS-509744 [70], providing strong evidence that Gads tyrosine phosphorylation in this *in vitro* assay can be attributed directly to Itk (Figure S3B).

**Figure 4.**
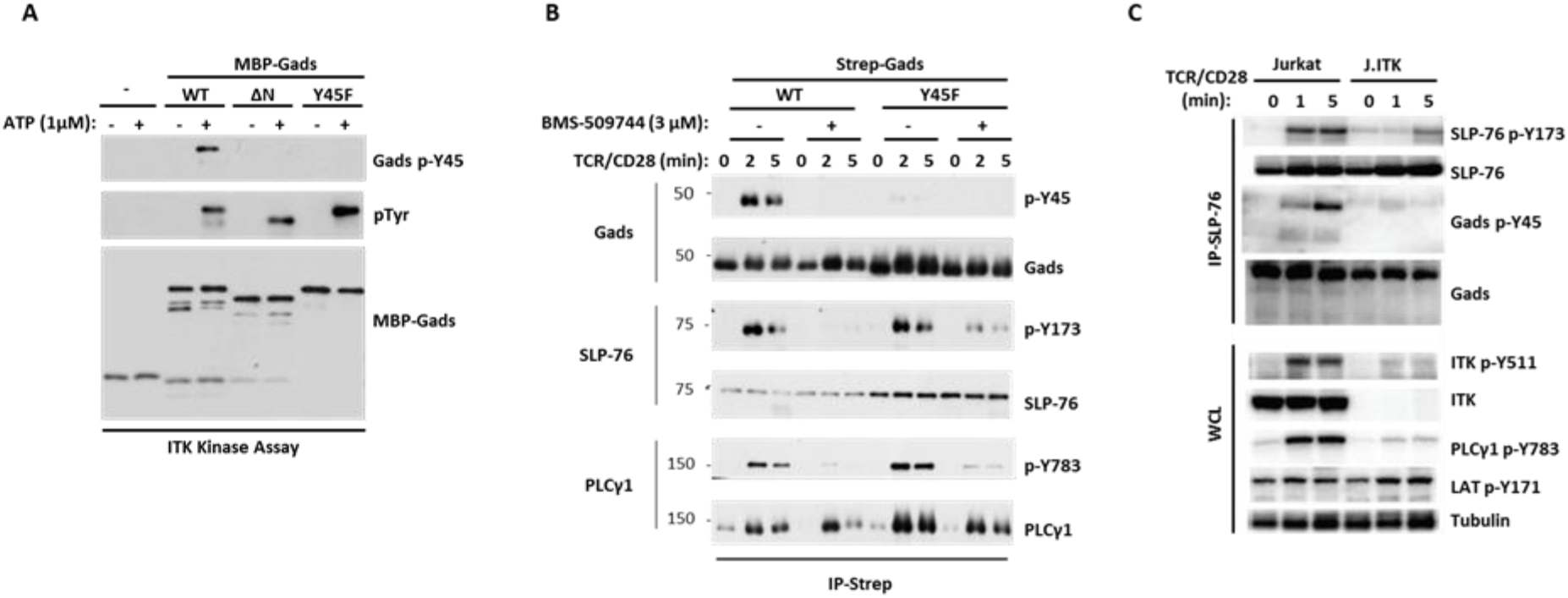
Itk mediates the TCR-inducible phosphorylation of Gads Y45 and SLP-76 Y173. (**A**) ***in vitro* phosphorylation of Gads Y45 by Itk.** Recombinant Gads, either full length (WT), lacking the N-terminal SH3 (ΔN) or with a substitution of phenylalanine for Tyr45 (Y45F), was phosphorylated *in vitro* by bead-bound Itk from TCR-stimulated cells, and the reaction supernatant was analyzed by blotting with anti-global p-Tyr (4G10), anti-p-Y45 or anti-MBP-Gads. (**B**) **An Itk inhibitor blocks TCR-induced Gads Y45 phosphorylation in intact cells.** dG32 cells, stably reconstituted with twin strep-tagged Gads, either WT or Y45F, were preincubated in stimulation medium for 30 min at 37°C, in the presence or absence of 3 μM BMS-509744, and were then stimulated with anti-TCR (C305, 1:1000 dilution) + CD28 costimulation (CD28.2 2 μg/ml). Streptactin complexes were probed with the indicated antibodies, as in Figure 3A. (**C**) **Gads Y45 phosphorylation is impaired in an ITK-deficient T cell line.** Jurkat or J.ITK cells were stimulated for the indicated time with anti-TCR + anti-CD28 and lysed. Anti-SLP-76 immune complexes and lysates were probed with the indicated phospho-specific antibodies, and then stripped, and reprobed for the total protein levels.

Our *in vitro* results provided evidence that Itk can directly phosphorylate Gads Y45, as well as an additional site or sites on Gads. In an effort to identify additional Itk-targeted sites, we performed a mass spectromic analysis of phospho-Gads from our *in vitro* assay system. This analysis identified Gads p-Y45 with high confidence, and also identified Gads phosphorylation at Y324 (data not shown). However, Gads p-Y324 was not detected in our SILAC-based mass spectrometry study of TCR-stimulated cells (Figure S1A), nor was it detected in other high throughput phospho-MS studies of TCR-stimulated cells, which did identify Gads Y45 [59, 60]. In the absence of evidence that Gads Y324 is phosphorylated in intact cells, we focused our attention on Gads Y45.

We employed pharmacologic and genetic approaches to test whether Itk mediates Gads Y45 phosphorylation in the context of intact T cells. In one approach, dG32 cells reconstituted with twin-strep-tagged Gads were stimulated in the presence of BMS-509744, a selective Itk inhibitor [70], and strep-tactin-purified Gads complexes were probed by immuno-blotting. BMS-509744 inhibited the TCR/CD28-induced phosphorylation of SLP-76 Y173 and PLC-γ1 Y783, both known Itk substrates, and likewise inhibited the phosphorylation of Gads Y45 (Figure 4B). As a control for specificity, we note that BMS-509744 did not inhibit the TCR-induced association of Gads with PLC-γ1, which is mediated by their mutual association with phospho-LAT (Figure 4B, bottom panel). In a complementary approach, phosphorylation of Gads Y45, SLP-76 Y173 and PLC-γ1 Y783 were abrogated in an Itk-deficient derivative of the Jurkat T cell line, J.ITK; as a control for specificity, the TCR-inducible phosphorylation of LAT Y171 was unaffected (Figure 4C).

Taken together, these results indicate that Gads Y45, like SLP-76 Y173 and PLC-γ1 Y783, is a *bona-fide* Itk substrate that is phosphorylated in intact cells in response to TCR signaling.

#### 3.3. SLP-76 targets active Itk to Gads Y45

Specific docking interactions are generally required to target Itk catalytic activity to its substrates [22, 46, 47]. Since active Itk associates with SLP-76 [20], and SLP-76 binds constitutively to Gads [71–73], we reasoned that SLP-76 may bridge the docking of catalytically active Itk onto its substrate, Gads Y45 (see Figure 1, right).

To test this idea, we reconstituted the Gads deficient cell line, dG32, with twin-strep-tagged Gads, either wild-type or P321L, a mutant form of Gads that does not bind to SLP-76 [53]. As expected, this mutation eliminated the constitutive association of SLP-76 with Gads (Figure 5A, panels 2 and 3). The Gads P321L mutation did not abolish Itk activation, as measured by its ability to phosphorylate SLP-76 at Y173; however, PLC-γ1 phosphorylation was substantially reduced (Figure 5A, bottom panels). These results are consistent with our previous observations in Gads-deficient T cells, and further support the notion that Gads is not required for TCR-mediated activation of Itk, but facilitates the Itk-mediated phosphorylation of PLC-γ1 by bringing SLP-76 associated Itk into the vicinity of LAT-associated PLC-γ1 [38].

**Figure 5.**
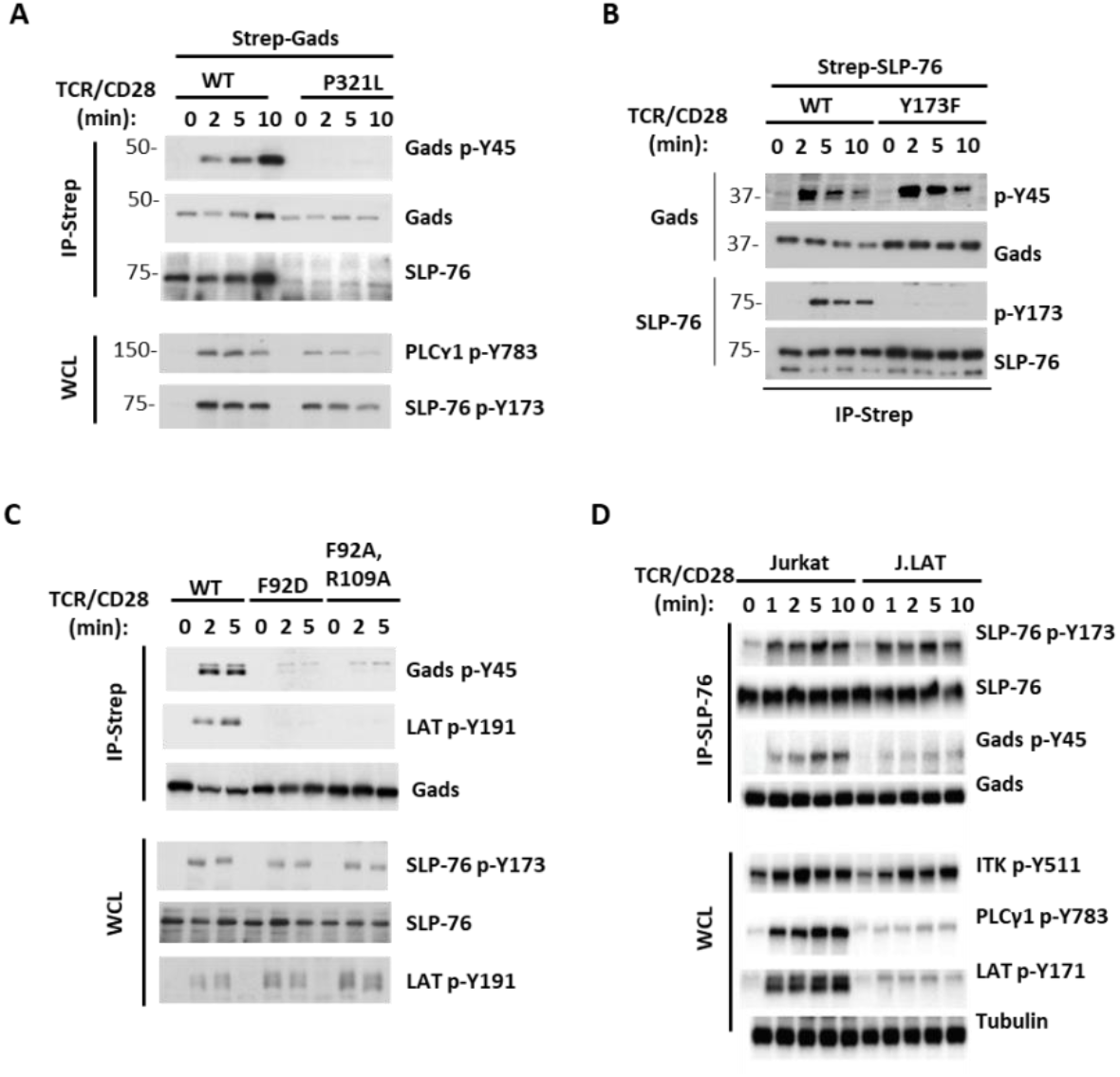
Gads Y45 phosphorylation occurs within the SLP-76-Gads-LAT complex. Jurkat-derived cell lines bearing specific mutations, as described in Table 1, were used to determine the structural requirements for Gads Y45 phosphorylation. Cells were stimulated for the indicated times with anti-TCR (C305, 1:1000 dilution) + CD28 costimulation (CD28.2 2 μg/ml) and lysed. Whole cell lysates (WCL) or affinity-purified complexes were probed with the indicated phospho-specific or total protein antibodies. (**A**) **The Gads-SLP-76 interaction targets Itk activity to Gads.** phospho-sites in dG32 cells, stably reconstituted with twin-strep-tagged Gads, either WT or bearing the C-SH3-inactivating mutation, P321L. (**B**) **Gads Y45 phosphorylation is independent of SLP-76 Y173.** Phospho-sites in J14 cells, stably reconstituted with twin strep-tagged SLP-76, either wild type or bearing the Y173F mutation. (**C**) **Gads dimerization is required for Y45 phosphorylation.** Phospho-sites were assessed in previously-described dG32 cell lines, stably reconstituted with WT or dimerization-deficient forms of Gads-GFP. (**D**) **LAT is required to support Gads Y45 phosphorylation.** TCR/CD28-induced phosphorylation events were compared in Jurkat, or the LAT-deficient cell line, J.LAT.

Whereas the Gads P321L mutation only partially reduced the phosphorylation of SLP-76 and PLC-γ1, it eliminated detectable phosphorylation of Gads Y45 (Figure 5A, top panel). Thus, the constitutive binding of SLP-76 to Gads is required in order to target SLP-76-bound Itk to Gads. It is important to emphasize that this requirement does not imply any type of hierarchical, or ordered phosphorylation of the two Itk-mediated sites; indeed, several of our results, presented above, clearly demonstrated that Gads p-Y45 is not required for the phosphorylation of SLP-76 Y173 (Figure 3A, 4B and 5A). Conversely, SLP-76 Y173 was not required for the TCR-induced phosphorylation of Gads Y45 (Figure 5B). These results suggest that within the SLP-76-Gads complex, SLP-76-bound Itk can be independently targeted to two different substrates, SLP-76 Y173 and Gads Y45.

#### 3.4. Gads Y45 phosphorylation occurs within the SLP-76-Gads-LAT complex

Next, we wondered whether the Gads-mediated recruitment of SLP-76 into the LAT-nucleated complex is required for Itk-mediated phosphorylation events. We disrupted the bridging activity of Gads by using two previously described mutant forms of Gads (Gads F92D, and Gads F92A,R109A), in which targeted disruption of the Gads SH2 dimerization interface disrupts its cooperatively-paired binding to LAT [Figure 5C and 42]. Whereas phosphorylation of SLP-76 Y173 was moderately reduced upon disruption of the Gads-LAT interaction; Gads Y45 phosphorylation was undetectable (Figure 5C). These results suggest that the SH2-mediated binding of Gads to LAT is required to direct Itk activity to Gads Y45.

To further validate this result, we disrupted LAT complex formation by CRISPR-mediated deletion of LAT. As expected, deletion of LAT abrogated the TCR-inducible phosphorylation of PLC-γ1; moreover, deletion of LAT greatly diminished the phosphorylation of Gads Y45, while only mildly reducing the phosphorylation of Itk Y511 and SLP-76 Y173 (Figure 5D).

Taken together, these experiments provide evidence for the existence of mechanistically independent pathways by which Itk activity is directed to its substrate sites on Gads and SLP-76. Gads Y45 phosphorylation absolutely depends on the interaction of Gads with both SLP-76 and LAT. In contrast, the TCR-inducible phosphorylation of SLP-76 Y173 is promoted by the binding of Gads to LAT but can occur in the absence of these adaptors.

#### 3.5. Distinct docking interactions direct Itk activity to its substrates, Gads, SLP-76 and PLC-γ1

We next turned our attention to the molecular mechanisms by which SLP-76-bound Itk is targeted to its different substrates. The SH2 and SH3 domains of Itk can mediate multiple, relatively weak interactions with SLP-76 [28, 74], and perhaps also with SLP-76-associated Vav [75]. This multiplicity of Itk-binding sites may allow for different conformations of SLP-76-bound Itk, each of which may be competent to phosphorylate a different substrate. To test this notion, we examined the role of two Itk-binding sites in mediating the phosphorylation of different Itk substrates.

Based on previous work [21, 30], we expected that the SH2-mediated binding of Itk to SLP-76 p-Y145 would be critical for phosphorylation of its substrates. To test this assumption, prior to the development of our p-Y45 phospho-specific reagent, we used a SILAC approach to quantitatively measure the TCR-induced fold-increase in Gads phosphorylation sites in TCR-stimulated J14 cells expressing twin-strep-tagged SLP-76 Y145F. We were surprised to observe pronounced TCR-induced phosphorylation of Gads Y45 (Figure 6A), which closely resembled our previous observations in WT SLP-76-expressing cells (Figure 2A).

**Figure 6.**
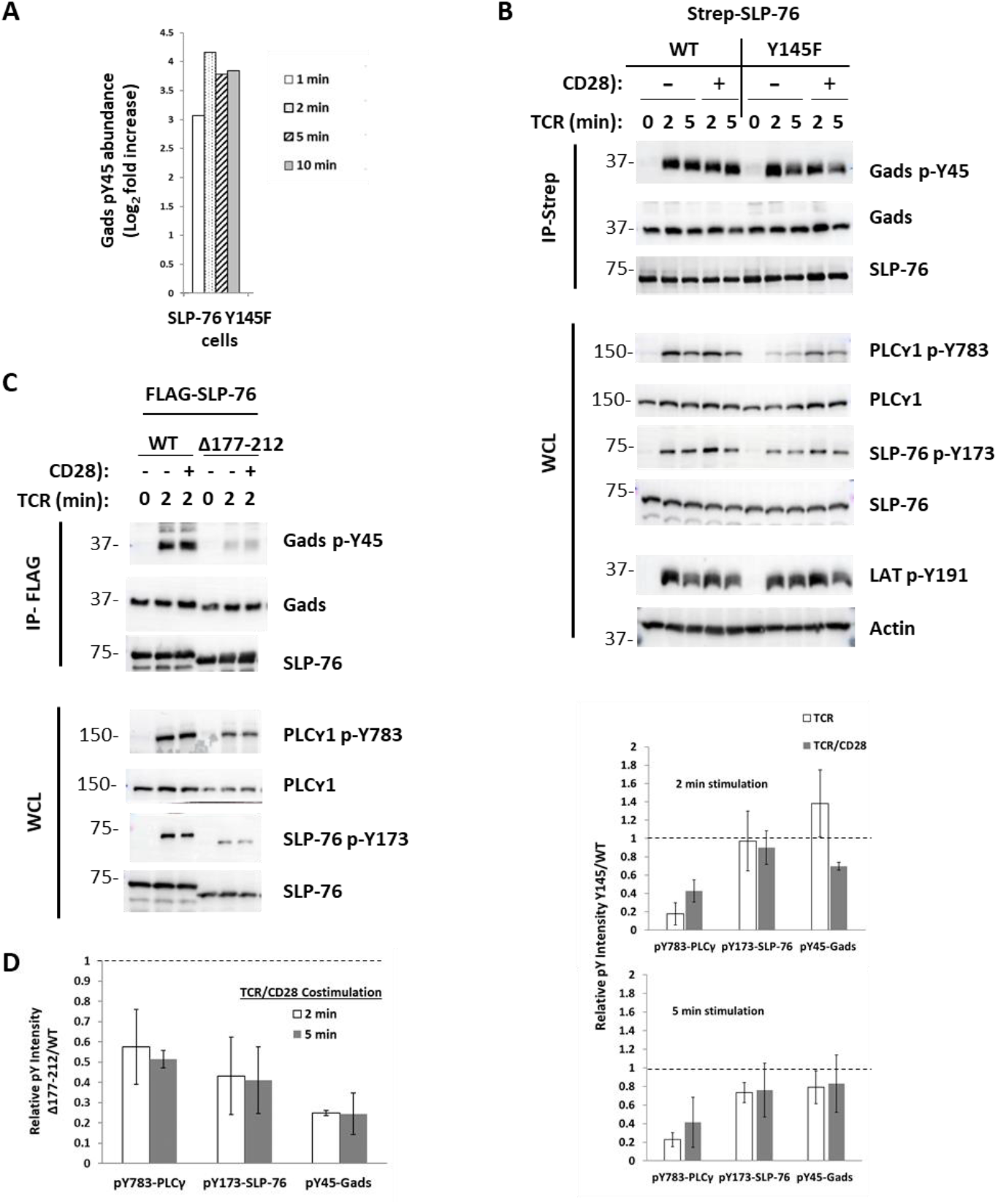
Distinct docking interactions direct Itk activity to its substrates, Gads, SLP-76 and PLC-γ1. J14 cells, stably reconstituted with the indicated forms of N-terminally strep-tagged (**A-B**) or FLAG-tagged (**C**) SLP-76, were stimulated as indicated and affinity-purified SLP-76 complexes or whole cell lysates (WCL) were analyzed to determine site-specific phosphorylation. (**A**) **SLP-76 Y145, an Itk SH2-binding site, is not required for Gads Y45 phosphorylation.** A SILAC approach was employed, as in Figure 2A, to identify the TCR-induced change in Gads Y45 phosphorylation in J14 cells expressing twin strep-tagged SLP-76 Y145F. Results are the median of two independent biological repeats. (**B**) **Differential effects of SLP-76 Y145 on Itk substrates.** TCR- and TCR/CD28-induced phospho-sites in J14 cells, stably reconstituted with SLP-76 WT or Y145F. Top: representative western blots. Bottom: Quantification of phospho-intensity. Gads pY45 phospho-intensity was normalized to total Gads protein. Results are expressed as the average phospho-intensity observed in SLP-76 Y145F-expressing cells, relative to the intensity observed in WT cells from the same stimulation time. n= 2 (TCR alone) or 3 (TCR/CD28) experiments, error bars represent the standard deviation. (**C**) **Differential effect of SLP-76 QQPP motif on Itk substrates.** TCR/CD28-induced phospho-sites in J14 cells, stably reconstituted with SLP-76 WT or Δ177-212. (**D**) **Quantitative effect of QQPP motif on Itk substrates.** The cell lines shown in C were stimulated for 2 or 5 minutes with TCR/CD28. Anti-FLAG complexes were probed for phospho- and total Gads, whereas lysates were probed for phospho-SLP-76 and phospho-PLC-γ1, and phospho-intensity was quantified as in B. n= 2 (5 min) or 3 (2 min) experiments, error bars represent the standard deviation.

We later recapitulated this surprising result by directly comparing SLP-76 WT and Y145F-expressing cells in immunoblotting experiments. Consistent with previous reports [21, 30], disruption of the Itk-binding site at SLP-76 Y145 markedly and consistently reduced the TCR-induced phosphorylation of PLC-γ1, although this effect was partially blunted by the addition of CD28 co-stimulation (Figure 6B, WCL panels and quantitation in bottom panels). In contrast, the SLP-76 Y145F mutation did not markedly reduce the TCR-induced phosphorylation of SLP-76 Y173 (Figure 6B WCL panels) or Gads Y45 (Figure 6B IP panels), either in the absence or in the presence of costimulation (see quantitation in bottom panels). These results provide evidence that different conformational states of SLP-76-bound Itk may be required to mediate its phosphorylation of PLC-γ1, as compared to SLP-76 and Gads.

To further explore this hypothesis, we examined the role of the QQPP motif, found at SLP-76 residues 184-195, which can serve as a ligand for the SH3 domains of Itk or of PLC-γ1 [28, 29, 32]. J14 cells were stably reconstituted with FLAG-tagged SLP-76, either WT or bearing a 36 amino acid deletion (Δ177-212) that encompasses the QQPP motif. Whereas the Δ177-212 deletion moderately reduced PLC-γ1 p-Y783 and SLP-76 p-Y173, Gads p-Y45 was profoundly reduced (Figure 6C and 6D). Thus, the QQPP Itk SH3 domain-binding motif appears to be important for Itk activation in general but is particularly required to direct Itk activity to Gads Y45.

Taken together, these experiments provide evidence that distinct conformations of SLP-76-bound active Itk may be required to direct its activity to particular substrates. A Y145-ligated conformation may primarily facilitate the phosphorylation of PLC-γ1 Y783 but not Gads Y45, whereas a QQPP-ligated conformation may be required to direct Itk activity to Gads Y45.

#### 3.6. Gads Y45 is not essential for TCR-proximal signaling to PLC-γ1

Having established that SLP-76 p-Y173 and Gads p-Y45 are TCR-inducible Itk-mediated phosphorylation sites within the LAT-nucleated complex, we next considered what the functional significance of these sites might be. Since Itk, SLP-76 and Gads are all implicated in the phosphorylation and activation of PLC-γ1 [20, 36, 38, 54, 76], we first explored the possibility that Gads p-Y45 may play a role in regulating TCR signaling to PLC-γ1.

To this end, we compared TCR-proximal signaling events in Gads WT- and Y45F-reconstituted dG32 cells. The Y45F mutation did not interfere with the TCR/CD28-induced association of Gads with phospho-LAT or with its indirect, LAT-mediated association with PLC-γ1 (Figure 7A, IP panels). The TCR-inducible phosphorylation of SLP-76 Y173 and LAT Y191 were likewise not affected by the Gads Y45F mutation (Figure 3A and 7A). Notably, phosphorylation of PLC-γ1 Y783 was unaffected, both within strep-tactin-purified Gads complexes, and in whole cell lysates (Figure 7A), suggesting that PLC-γ1 recruitment to LAT, phosphorylation and its release from the LAT complex all proceed independently of Gads Y45.

**Figure 7.**
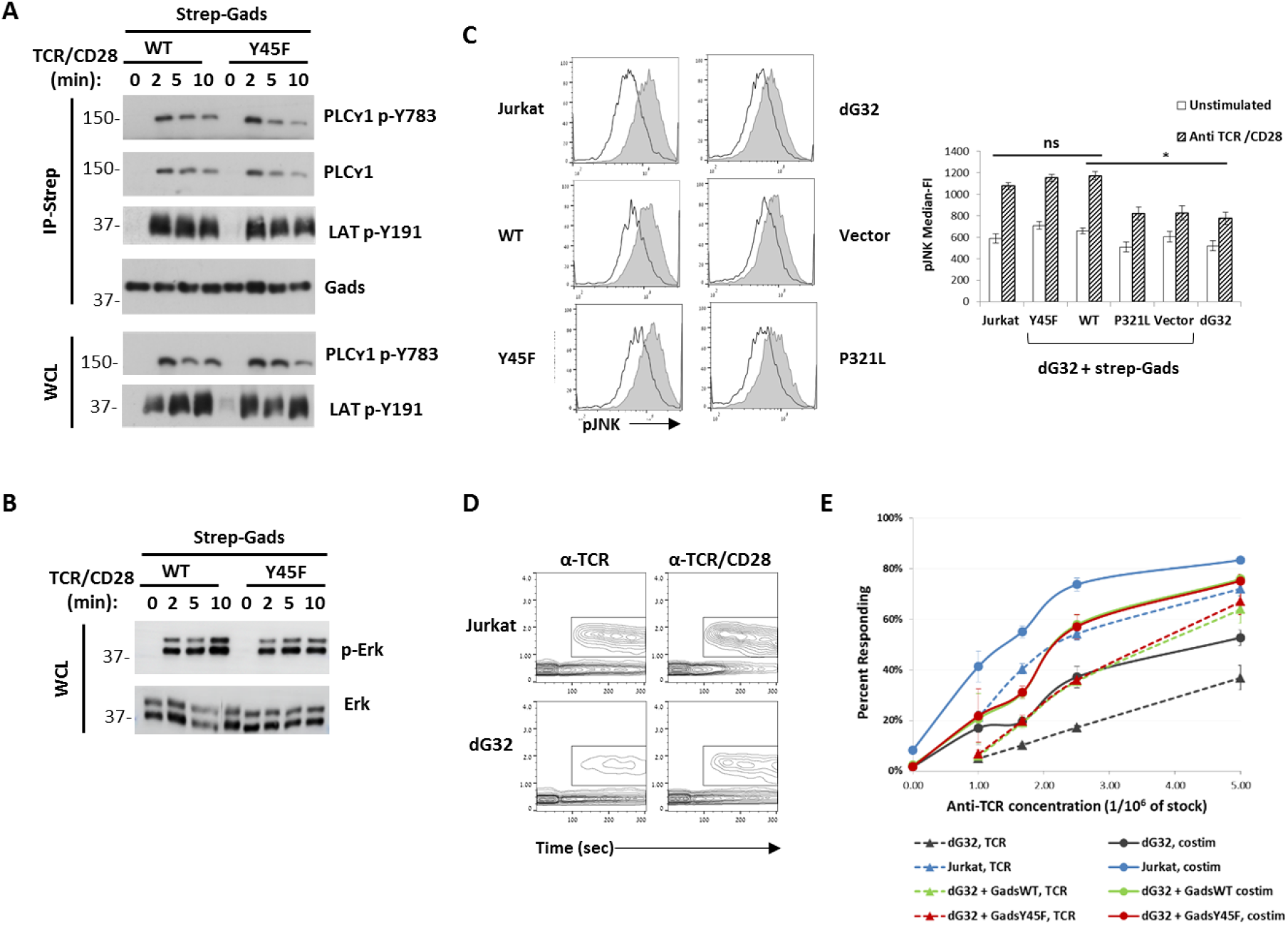
Gads Y45 is not required for TCR-proximal signaling to PLC-γ1. (**A**) **Recruitment of Gads to LAT is independent of Gads p-Y45.** Lysates (WCL), or streptactin-purified Gads complexes from stimulated cells were probed with the indicated antibodies. Results are representative of at least 3 repeats. (**B**) **Erk MAPK activation is independent of Gads pY45.** Cells were stimulated as in A and lysates were probed with anti-phospho- and total Erk1/2. (**C-E**) FACS-based assays were used to measure TCR responses in Jurkat, dG32 or dG32 cells that were stably reconstituted with the indicated twin strep-tagged Gads alleles. Cells were differentially barcoded with CellTrace Violet (**C**) or CellTrace Far Red (**D-E**), prior to stimulation. (**C**) **pJNK response is independent of Gads p-Y45.** Cells were stimulated in triplicate for 15 min with anti-TCR (1:30,000) + anti-CD28 (2 μg/ml) (shaded histogram), or mock stimulated (open histogram), prior to intracellular staining with anti-p-JNK-PE. Results were analyzed while gating on matched GFP-expression gates, used as an indication of Gads expression. Left: Representative results. Right: Average p-JNK median fluorescence intensity (n=3; error bars indicate the SD). The unpaired two-tailed T test was used to identify statistically significant differences, relative to TCR-stimulated WT-reconstituted dG32 cells (*, p<0.05). (**D-E**) **Calcium response is independent of Gads p-Y45.** Intracellular calcium was measured by FACS, with TCR or TCR/CD28 costimulation added at 60 sec. (**D**) Representative raw data observed upon stimulation with anti-TCR (C305 1:400,000), in the presence or absence of anti-CD28 (1.5 μg/ml). Cells within the rectangular gate are considered to be responding cells. (**E**) Cells were stimulated with the indicated concentration of anti-TCR (C305), in the presence or absence of CD28 costimulation (1.5 μg/ml). Shown is the percent responding cells observed within a 100 second time window, beginning 1.5 min after the addition of stimulant (n=3, error bars indicate SD).

Further downstream, the Gads Y45F mutation did not reduce the TCR/CD28-induced phosphorylation of MAPK family members, Erk (Figure 7B) or Jnk (Figure 7C), nor did it mitigate the TCR-induced increase in CD69 expression (Figure S4A).

To more formally rule out a role for Gads Y45 in regulating PLC-γ1 activation, we examined its effect on the TCR-induced calcium response, by assessing the frequency of cells that exhibited increased intracellular calcium in response to low dose TCR stimulation. As we previously reported [38], Gads-deficiency markedly reduced the frequency of responding cells (Figure 7D, left) over a range of TCR stimulatory doses (Figure 7E). Stable reconstitution of Gads expression increased the sensitivity of TCR responsiveness in this assay; however, the magnitude of the Gads-dependent increase was not affected by the Y45F mutation (Figure 7E, dotted lines). CD28 costimulation further increased the frequency of responding cells, both in the presence and in the absence of Gads (Figure 7D); moreover, the CD28-dependent increase in responsiveness was not affected by the Gads Y45F mutation, as compared to WT-reconstituted cells (Figure 7E, solid lines).

Taken together, these results provide evidence that Gads p-Y45 is not required for TCR signaling to PLC-γ1.

#### 3.7. SLP-76 Y173 exerts a modest effect on TCR-proximal signaling to PLC-γ1

Having excluded a role for Gads p-Y45 in mediating PLC-γ1 activation, we decided to take a closer look at the role of SLP-76 p-Y173. Our previously published data suggested, but did not definitively prove a role for SLP-76 Y173 in regulating PLC-γ1. Whereas the Y173F mutation decreased the TCR-induced accumulation of phospho-PLC-γ1 in whole cell lysates; this mutation only modestly reduced TCR-induced calcium flux [21]. To address this apparent contradiction, we compared the accumulation of PLC-γ1 p-Y783 within two pools of PLC-γ1, the SLP-76-bound pool, and the pool that is found in whole cell lysates. The latter pool includes PLC-γ1 that was phosphorylated within the LAT-nucleated complex and subsequently released [64]. Whereas the Y173F mutation markedly reduced the abundance of PLC-γ1 p-Y783 in whole cell lysates, the SLP-76-bound pool of PLC-γ1 p-Y783 was not reduced (Figure S4B). This observation suggests that SLP-76 Y173 is not required for PLC-γ1 phosphorylation *per se*, but may be required for the release of phosphorylated PLC-γ1 from the LAT-nucleated complex into the cytosol.

#### 3.8. The Itk-targeted sites on SLP-76 and Gads are selectively required for TCR/CD28 signaling to the RE/AP transcriptional element

Having demonstrated that PLC-γ1 activation occurs independently of Gads Y45, and is partially independent of SLP-76 Y173, we considered the possibility that the Itk-targeted sites on SLP-76 and Gads may regulate a distinct aspect of the TCR/CD28 signaling pathway.

The LAT-nucleated complex controls different branches of the TCR signaling pathway, leading to the activation of different transcriptional elements, which together drive the transcription of IL-2 [1]. PLC-γ1 produces two second messengers, IP3 and DAG, which respectively bring about the activation of NFAT and AP1 transcription factors that bind to a compound NFAT/AP1 site within the IL-2 promoter. A second compound site, RE/AP, binds to AP1 and NFκB, and is activated in response to TCR/CD28 costimulation [7].

Whereas SLP-76 is absolutely required for NFAT and AP-1 activation [54], the Y173F mutation did not reproducibly affect the activity of an NFAT/AP1 luciferase reporter construct (Figure 8A, left). Nevertheless, this mutation markedly reduced the TCR/CD28-induced production of IL-2 (Figure 8B), suggesting that it may affect signaling through a different branch of the TCR signaling pathway. Consistent with this notion, the Y173F mutation eliminated TCR/CD28-induced activation of an RE/AP-luciferase reporter construct (Figure 8A, middle) and markedly reduced the activation of an NFκB-luciferase reporter construct (Figure 8A, right).

**Figure 8.**
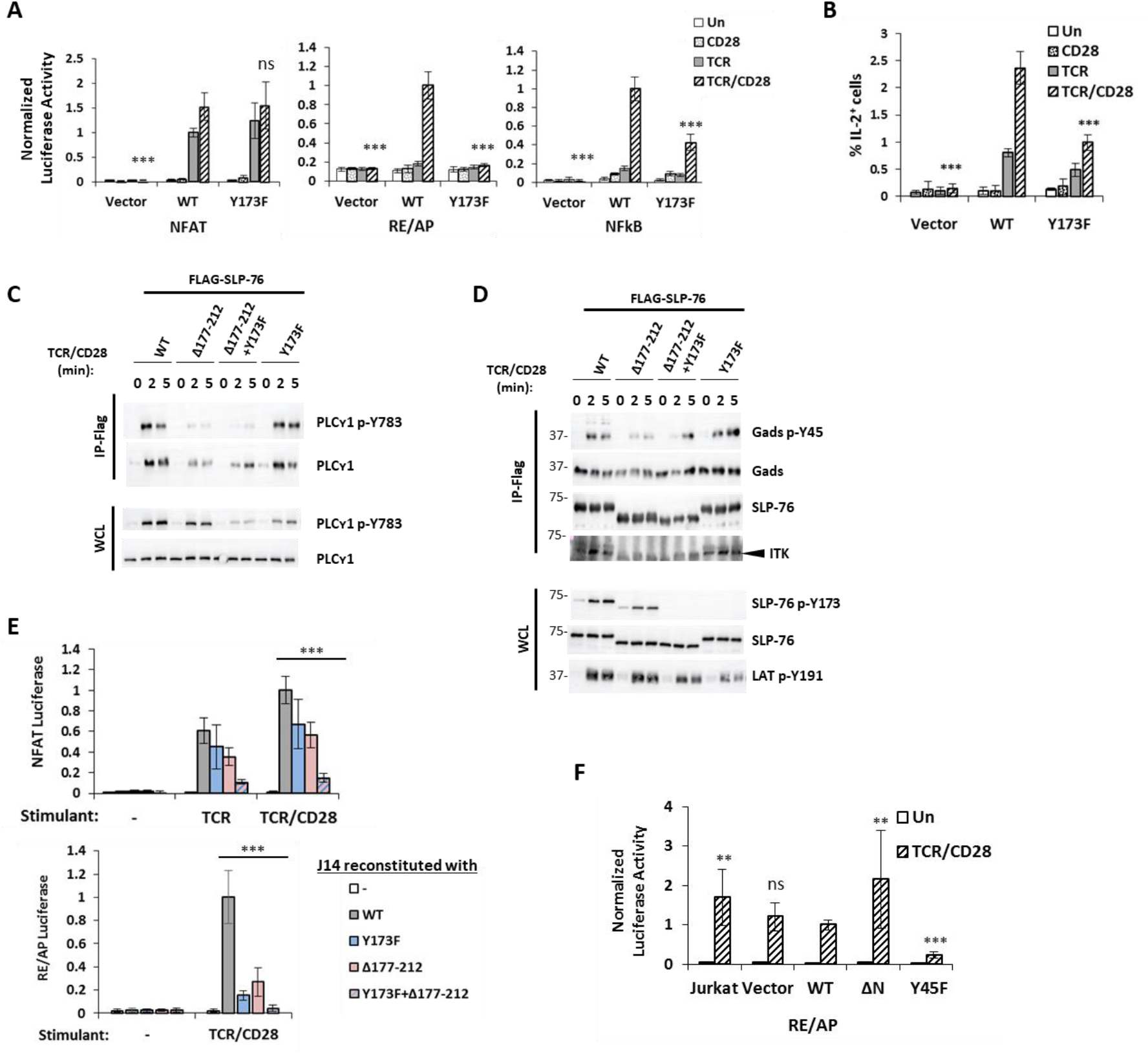
Itk-related features of SLP-76 and Gads are specifically required to activate the RE/AP transcriptional element. (**A**) **Role of SLP-76 Y173 in transcriptional responses to TCR/CD28 stimulation.** Luciferase reporter activity was measured in J14 cells that had been stably reconstituted with comparable expression of twin strep-tagged SLP-76 (WT or Y173F) or with a vector control. Normalized results are presented relative to that observed in TCR/CD28-stimulated WT cells from the same stimulation plate. Results are the average of 3 experiments, conducted in triplicate, error bars indicate the SD. (**B**) **Role of SLP-76 Y173 in TCR/CD28-induced IL-2 expression.** The cell lines shown in A were stimulated for 6 hours with plate-bound anti-TCR and soluble anti-CD28, in the presence of 5 μg/ml brefeldin A during the last 4 hours of stimulation, and intracellular staining with α-IL-2-PE was analyzed by FACS, to determine the percent IL-2^+^ cells. Results are the average of 2 experiments, conducted in duplicate, error bars indicate the SD. (**C-E**) **Two Itk-related features of SLP-76 are required for RE/AP activation.** J14 cells were reconstituted with the indicated forms of FLAG-tagged SLP-76 and were sorted for comparable expression level. (**C-D**) **Differential effect of the SLP-76 mutations on PLC-γ1 and Gads phosphorylation.** Cells were stimulated and western blots prepared from anti-FLAG-purified SLP-76 complexes (IP-FLAG) or whole cell lysates (WCL). (**E**) **NFAT and RE/AP luciferase activities** were measured as in A. Results are the average of 3 or 4 experiments, conducted in triplicate, error bars indicate the SD. (**F**) **Role of Gads Y45 in transcriptional response to TCR/CD28.** dG32 cells were stably reconstituted with twin-strep tagged Gads (WT, ΔN or Y45F) or with a vector control and were sorted for comparable expression level. RE/AP luciferase activity was measured as in A. Results are the average of four experiments, each conducted in triplicate, error bars indicate the SD. For all bar graphs in this figure, the unpaired student T test was used to compare TCR/CD28-stimulated cells to WT TCR/CD28-stimulated cells (**, p<0.005; ***, p<0.0005; ns, p>0.05).

These results suggest that SLP-76 p-Y173 is selectively required for an NFκB-dependent branch of the signaling pathway in which TCR and CD28 costimulation synergistically bring about the activation of RE/AP. Moreover, these results imply that Itk may exert two mechanistically distinct influences on IL-2 transcription. Via phosphorylation of PLC-γ1, Itk promotes activation of the NFAT transcriptional element, whereas via phosphorylation of SLP-76 Y173, Itk promotes the activation of RE/AP.

To further explore the dual role of Itk as a regulator of NFAT and RE/AP, we chose to compare two mutations that disrupt Itk-mediated signaling in different ways. To this end, we reconstituted J14 cells with FLAG-tagged SLP-76, either WT, Y173F, Δ177-212 (which lacks the QQPP motif), or with an allele of SLP-76 bearing both mutations, Δ177-212+Y173F. This setup allowed us to compare different effects of Itk: whereas the Y173F mutation disrupts Itk-mediated phosphorylation of SLP-76, the Δ177-212 mutation primarily disrupts Itk-mediated phosphorylation of Gads Y45, and the double mutation disrupts the phosphorylation of SLP-76, Gads and PLC-γ1 (Figure 8C and 8D)

Activation of NFAT was only modestly reduced by each of the single mutations, Y173F or Δ177-212, but was dramatically reduced in cells bearing the double mutation, SLP-76 Y173F+Δ177-212 (Figure 8E, top). The inhibitory effect of the combined mutations may be understood in terms of their effects on PLC-γ1 phosphorylation (Figure 8C). The Y173F mutation decreased PLC-γ1 p-Y783 in whole cell lysates, but not within the SLP-76-bound pool. The Δ177-212 mutation had the opposite effect, dramatically decreasing PLC-γ1 p-Y783 within the SLP-76-bound pool, but only moderately decreasing PLC-γ1 phosphorylation in whole cell lysates. The double mutation synergistically impaired NFAT activity, most likely due to its ability to markedly reduce the phosphorylation of PLC-γ1 both in both compartments.

Compared with their modest effect on NFAT activation (Figure 8E, top), each of the single mutations markedly impaired the activation of RE/AP, which was even more profoundly impaired by the double mutation (Figure 8E, bottom). These results suggest the presence of at least two PLC-γ1-independent mechanisms by which Itk can influence downstream signaling events leading to RE/AP. First, Itk is required for SLP-76 Y173 phosphorylation, which is required to support RE/AP activation. Second, Itk may act via the QQPP motif to influence RE/AP activation by an additional mechanism. While neither mechanism on its own is required for NFAT activation, removal of both mechanisms abrogated NFAT activity as well as RE/AP activity. Since the Δ177-212 mutation dramatically decreased Gads p-Y45 but only moderately affected SLP-76 p-Y173, we considered the possibility that Gads p-Y45 may be required for RE/AP activation.

Gads plays a supporting role in the TCR signaling pathway, for example, it promotes, but is not absolutely required for TCR-induced calcium flux and NFAT activation [38]; moreover, the thymic developmental defect of Gads-deficient mice is much milder than the absolute thymic block exhibited by SLP-76-deficient mice [3]. Consistent with this notion, TCR/CD28-induced RE/AP activation was Gads-independent. Nevertheless, the Y45F mutation dramatically reduced activation of the RE/AP reporter (Figure 8F). To verify this surprising result, we independently repeated the reconstitution of dG32 cells, and again found that whereas RE/AP activation can occur equally well in the absence or presence of Gads, reconstitution of the cells with Gads bearing a mutation at Y45, either Y45F or Y45E, resulted in profound inhibition of RE/AP activity (Figure S4B).

This unexpected result suggested that Gads may perform an inhibitory signaling function, which must be removed in order to activate RE/AP. According to this model, the inhibitory function may be removed upon TCR-induced phosphorylation of Gads Y45. Consistent with this result, removal of the N-SH3 did not impair RE/AP activation, and in some experiments activation was modestly increased (Figure 8F and Figure S4B).

Taken together, these results suggest that Itk controls CD28 responsiveness via TCR-induced phosphorylation of its targets on Gads and SLP-76. Moreover CD28 responsiveness appears to be actively restrained by un-phosphorylated Gads. The fact that two independent pathways are required for RE/AP activation - a SLP-76-dependent pathway leading to SLP-76 Y173 phosphorylation and a LAT-dependent pathway leading to Gads Y45 phosphorylation - may help to insure the interdependence of CD28 responsiveness on TCR signaling.

## 4. Discussion

Prior studies established a clear role for Itk in the initiation of TCR-proximal signaling events, leading to the phosphorylation and activation of PLC-γ1 [25]. As a Tec-family kinase that is activated downstream of Lck and ZAP-70, Itk constitutes the third member of the TCR-induced tyrosine kinase cascade. Active Itk can be found in complex with SLP-76, which is recruited to LAT by Gads, and within this heterotrimeric adaptor complex, Itk promotes downstream responsiveness, in part, by phosphorylating LAT-associated PLC-γ1 at Y783.

Here, we have uncovered a distinct Itk-dependent signaling module that is based on the Itk-mediated phosphorylation of SLP-76 Y173 and Gads Y45. Our data document the profound TCR-induced increase in the abundance of SLP-76 p-Y173 and Gads p-Y45, both in a human T cell line and in primary mouse lymphocytes [Figure 2-3 and 21]. We provide several independent lines of evidence to demonstrate that these phosphorylation events are mediated by Itk (Figure 4), and that each site can be phosphorylated independently of the other (Figures 3A and 5B). Itk-mediated phosphorylation of Gads occurs within the LAT-nucleated signaling complex (Figure 5) and appears to depend on a particular conformation of Itk, in which the Itk SH3 domain binds to the QQPP motif of SLP-76 (Figure 6). Whereas Gads p-Y45 and SLP-76 p-Y173 are largely expendable for signaling through the PLC-γ1-calcium-NFAT axis; both are required to mediate the TCR/CD28-induced activation of the RE/AP transcriptional element from the IL-2 promoter (Figure 7-8). This study therefore establishes a PLC-γ1-independent mechanism by which SLP-76 and Gads regulate the synergistic activity of the TCR and CD28 signaling pathways, leading to the activation of RE/AP.

The high fold phosphorylation and evolutionary conservation of the Gads p-Y45 motif, which is found within the conserved N-terminal SH3 domain of Gads, all suggest a conserved function. Yet the N-terminal SH3 domain of Gads, to date, has no known ligand or signaling function [3], and our characterization of Gads p-Y45 therefore provides the first insight into the regulatory role played by the N-SH3 of Gads. Moreover, our study may provide some insight into the widespread but poorly understood phenomenon of SH3 domain tyrosine phosphorylation [77]. A recently published bioinformatic survey revealed that of 273 human SH3 domains, 94 are phosphorylated on tyrosine, and 20 of those are phosphorylated at the M2 position, which is the location of Gads Y45 [78]. Yet, with the possible exception of Src p-Y133 [79] and Caskin 1 p-Y336 [80], the functional significance of SH3 domain tyrosine phosphorylation at the M2 position is, to the best of our knowledge, unknown.

Upon the identification of Gads Y45 and SLP-76 Y173 as substrates of Itk, we sought to identify the mechanisms by which the catalytic domain of Itk is directed to these particular substrates. This question is complicated by the fact that Itk is regulated by multiple intra- and inter-molecular protein-protein interactions [25]. In the resting state, intramolecular interactions of the SH2 and SH3 domains of Itk stabilize the inactive conformation of the enzyme; whereas upon TCR stimulation, bivalent binding of SLP-76 to the SH domains of Itk is thought to stabilize the catalytically active conformation [24, 28]. Consistent with this notion, elution of Itk from the SLP-76 complex abrogated its activity, which was restored upon its reassociation with SLP-76 [20].

The association of active Itk with SLP-76 appears to be sufficient to target its activity to SLP-76 Y173. In support of this notion, we observed that the phosphorylation of SLP-76 at Y173 can occur in the absence of either Gads or LAT, and was only moderately reduced by point mutations that disrupt the Gads-mediated recruitment of SLP-76 to LAT [Figure 5 and 38].

In contrast, Itk-mediated phosphorylation of Gads was exquisitely dependent on the association of Gads with both SLP-76 and LAT, suggesting that Gads Y45 phosphorylation must occur within the heterotrimeric adaptor complex (Figure 5). These findings provide clues to the specific docking interactions that target Itk catalytic activity to Gads. In particular, the high affinity, constitutive interaction of Gads with SLP-76 [33, 40, 41] may serve to bring Gads Y45 into the vicinity of SLP-76-bound Itk. Moreover, the cooperatively paired binding of the Gads SH2 to LAT [42] may create a docking surface for the Itk kinase domain, which may facilitate the Itk-mediated phosphorylation of the adjacent Gads Y45. This speculation is consistent with previous studies, in which docking of the Itk kinase domain onto a non-classical SH2 domain surface within particular substrate proteins was required for the Itk-mediated phosphorylation of an SH2-adjacent site [46, 47]. Alternatively, the dimeric binding of Gads to LAT may induce a conformational change that may increase the surface exposure of Gads Y45. It is important to note that no structural information is available for the N-terminal SH3 of Gads, and we expect that future structural studies will be required to illuminate the mechanism by which Itk activity is targeted to Gads Y45.

To better understand the mechanisms that direct Itk to its individual substrates, we explored how phosphorylation of each substrate may depend on two known Itk-binding motifs: SLP-76 p-Y145, which binds to the SH2 domain of Itk, and the SLP-76 QQPP motif, which binds to the Itk SH3 domain (Figure 6). As previously reported [21, 30], ablation of SLP-76 Y145 markedly reduced the Itk-mediated phosphorylation of PLC-γ1; however, its effects on the Itk-mediated phosphorylation of SLP-76 and Gads were quite modest. Conversely, a deletion encompassing the QQPP motif markedly reduced the TCR-induced association of Itk with SLP-76 and the Itk-mediated phosphorylation of Gads Y45 but had a more modest effect on the phosphorylation of SLP-76 and PLC-γ1. These results provide evidence that Itk may assume multiple conformations within the SLP-76-Gads-LAT complex, each of which may be most suited to phosphorylation of a particular substrate within this complex.

The concept that Itk can assume distinct, catalytically active, SLP-76-bound conformations may help to resolve other previously puzzling observations. The SH2 of Itk is commonly thought to bind to SLP-76 pY145, yet ablation of SLP-76 Y145 was not sufficient to disrupt the interaction of Itk with SLP-76 [30]. The residual association of Itk with SLP-76 Y145F may be mediated by binding of the Itk SH3 to the QQPP motif, by binding of the Itk SH2 to pY113 of SLP-76, or by indirect recruitment of Itk, via its interaction with Vav [75], which binds directly to SLP-76 and may serve to stabilize the SLP-76-Itk interaction.

The binding site of the Itk SH3 domain is likewise not completely clear. Whereas it may bind to the QQPP motif [28, 29], this motif was also reported to bind to the SH3 domains of Lck and PLC-γ1 [32, 34]. One possible solution to this conundrum might involve the binding of the QQPP motif to different SH3 domains at different stages in the signaling cascade, or via partially overlapping binding sites, as was recently demonstrated for the binding of Itk and Lck to the related adaptor TSAD [81]. A switch between different QQPP-binding partners may be facilitated by the autophosphorylation of Itk at Y180 within its SH3 domain [66], as phosphorylation of this site alters the affinity of the SH3 for different proline-rich ligands [82]. In this way, autophosphorylation of Itk Y180 may alter its mode of binding to SLP-76.

As an integrative explanation for the above data, we propose that Itk may interact with SLP-76 via multiple distinct modes, each of which is most suitable for the phosphorylation of a particular substrate.

Further downstream, Gads Y45 and SLP-76 Y173 appear to be dispensable for the canonical SLP-76-Gads-LAT-mediated phosphorylation and activation of PLC-γ1, leading to calcium flux and NFAT activation; rather both sites are required to mediate the TCR/CD28-induced activation of the RE/AP transcriptional element (Figures 7-8). In a similar manner, a region encompassing the QQPP motif on SLP-76, was required for phosphorylation of Gads Y45 and for the activation of RE/AP but not NFAT (Figures 6C and 8C-E). Consistent with our findings, precise excision of the 10 amino acid QQPP motif in SLP-76-reconstituted J14 cells moderately reduced NFAT nuclear translocation, but dramatically reduced IL-2 production [34]. Our results therefore suggest that Itk regulates at least two distinct signaling branches downstream of the TCR, one leading to PLCγ1-dependent calcium flux, and the other acting via phosphorylation sites on SLP-76 and Gads to regulate the activity of the RE/AP transcriptional element.

Whereas we previously reported that SLP-76 p-Y173 is required for optimal phosphorylation of PLC-γ1 [21], a careful re-appraisal of this site revealed that SLP-76 p-Y173 is not required for PLC-γ1 phosphorylation *per se*, but rather contributes to the catalytic release of phospho-PLC from the LAT-nucleated complex [as described by 64], thereby promoting the accumulation of phospho-PLC-γ1 outside the confines of this complex (Supp Figure 4B). The ability of SLP-76 p-Y173 to promote the release of phospho-PLC-γ1 from LAT may relate to its ability to bind weakly to the C-terminal SH2 of PLC-γ1 [22]. It is interesting to note a previous mutational study of PLC-γ1, which presented evidence that the C-SH2 of PLC-γ1 is required for the activation of RE/AP, independently of any influence on TCR-induced calcium flux [83]. Thus, we can speculate that the pathway by which SLP-76 p-Y173 regulates RE/AP may be related to its ability to bind to the C-terminal SH2 of PLC-γ1.

The mechanisms by which Itk-mediated phosphorylation of Gads and SLP-76 regulate the activity of the RE/AP transcriptional element remain to be determined. One likely possibility is an effect on NFκB signaling, as revealed by the partial inhibition of an NFκB reporter upon mutation of SLP-76 Y173 (Figure 8A). Previous reports suggested that SLP-76 may regulate NFκB through HPK1, a kinase that associates with the SH2 domain of SLP-76 [84] and is reported to phosphorylate Carma1, a key element of the NFκB signaling pathway [85]. Another possibility is that SLP-76 and Gads may directly influence transcriptional events via the ability of SLP-76 to translocate to the nuclear pores, where it regulates the nuclear translocation NFκB [86]. It remains to be seen whether this activity depends on Itk-targeted phosphorylation sites on the adaptors. Another possibility is a direct effect on CD28 signaling. It has been known for some time that Gads can bind directly to a membrane-proximal pYMNM motif found in the cytoplasmic tail of CD28 [87]; however, evidence for the functional relevance of this interaction is mixed [88–90]. While it is possible that Gads p-Y45 may exert its effects directly within the CD28-nucleated signaling complex, we consider this possibility unlikely, since Gads Y45 did not affect the CD28-induced augmentation of calcium flux (Figure 7). In light of the large number of possibilities, a specific resolution of the mechanism by which SLP-76 p-Y173 and Gads p-Y45 regulate RE/AP is outside the scope of this study.

The profound inhibition of RE/AP by the Gads Y45F mutation is especially intriguing, as Gads itself was not required for RE/AP activation. This observation suggests that Gads performs an inhibitory function, analogous to the closing of a gate, which limits T cell responsiveness by inhibiting the activation of RE/AP. A similar phenomenon has been observed for the adaptor protein ALX, which is dispensable for RE/AP activation, but profoundly inhibits RE/AP when overexpressed [91, 92]. This example provides evidence for the existence of regulatory pathways that are dedicated to the negative regulation of RE/AP, and we suggest that Gads is likely to constitute an important component of this regulatory mechanism. The gating mechanism is likely to involve particular ligands of the N-SH3 of Gads, or a phospho-dependent ligand of Y45, which remain to be identified.

Based on this model, opening of the gate to allow RE/AP activation would depend on Gads SH2 domain dimerization, leading to the phosphorylation of Gads at Y45, which together may stabilize an active conformation. In this respect, it is important to note the tight regulation of Gads Y45 phosphorylation, induction of which depends both on the TCR-induced binding of Gads to LAT, and on the TCR-induced interaction of Itk with SLP-76. Since the gate remains closed and RE/AP is inhibited in the absence of Gads Y45 phosphorylation, these requirements may prevent spurious immune responses by limiting CD28 responsiveness to cells that have experienced signaling through the TCR.

## Supplementary Materials

Figure S1: Summary of SLP-76 and Gads phosphorylation sites identified in our SILAC-based phospho-mass spectrometry analysis, Figure S2: Evolutionary conservation of the Gads N- and C-terminal SH3 domains, Figure S3: Gads Y45 is an Itk-targeted site, Figure S4: Downstream signaling functions of Gads p-Y45 and SLP-76 p-173.

## Author Contributions

EH and DY developed the study objectives and experimental strategy with key inputs from HU, AW and W-LL. Mass spectrometry samples were prepared by DB and were analyzed by JC under the supervision of HU. Novel cell lines and immunological reagents used in this work were created by EH, RS, W-LL, IO, SW, AI, MS and DB. Cellular responses to stimulation were measured and data were analyzed by EH, RS, W-LL, IO, SW and DY. This manuscript was written by DY and EH, with key inputs from W-LL, AW and HU. All authors have read and agreed to the published version of the manuscript.

## Acknowledgments

The Biomedical Core Facility (BCF) of the Rappaport Faculty of Medicine provided access to FACS equipment, and BCF staff members Ofer Shenkar, Amir Grau, and Rotem Honen Kadosh provided excellent technical support. Gads deficient mice on the Balb/C background were generously provided by C. Jane McGlade (University of Toronto), We thank Dr. Rona Shofti from the Technion preclinical authority and her staff for their professional assistance with the care and housing of our mice.

## References

1. Gaud, G., R. Lesourne, and P.E. Love. 2018. Regulatory mechanisms in T cell receptor signalling. Nature Reviews Immunology.

2. Balagopalan, L., N.P. Coussens, E. Sherman, L.E. Samelson, and C.L. Sommers. 2010. The LAT Story: A Tale of Cooperativity, Coordination, and Choreography. Cold Spring Harbor Perspectives in Biology. 2:(8): p. a00 5512.

3. Yablonski, D. 2019. Bridging the Gap: Modulatory Roles of the Grb2-Family Adaptor, Gads, in Cellular and Allergic Immune Responses. Frontiers in Immunology. 10(1704): p. doi: 10.3389/fimmu.2019.01704.

4. Esensten, Jonathan H., Ynes A. Helou, G. Chopra, A. Weiss, and Jeffrey A. Bluestone. 2016. CD28 Costimulation: From Mechanism to Therapy. Immunity. 44(5): p. 973–988.

5. Tuosto, L. 2011. NF-κB family of transcription factors: Biochemical players of CD28 co-stimulation. Immunology Letters. 135(1): p. 1–9.

6. Fraser, J.D., B.A. Irving, G.R. Crabtree, and A. Weiss. 1991. Regulation of interleukin-2 gene enhancer activity by the T cell accessory molecule CD28. Science. 251(4991): p. 313–6.

7. Shapiro, V.S., K.E. Truitt, J.B. Imboden, and A. Weiss. 1997. CD28 mediates transcriptional upregulation of the interleukin-2 (IL-2) promoter through a composite element containing the CD28RE and NF-IL-2B AP-1 sites. Mol Cell Biol. 17(7): p. 4051–8.

8. Shapiro, V.S., M.N. Mollenauer, and A. Weiss. 1998. Nuclear factor of activated T cells and AP-1 are insufficient for IL-2 promoter activation: requirement for CD28 up-regulation of RE/AP. J Immunol. 161(12): p. 6455–8.

9. Matzinger, P. 2002. The Danger Model: A Renewed Sense of Self. Science. 296(5566): p. 301–305.

10. Brownlie, R.J. and R. Zamoyska. 2013. T cell receptor signalling networks: branched, diversified and bounded. Nat Rev Immunol. 13(4): p. 257–69.

11. Au-Yeung, B.B., N.H. Shah, L. Shen, and A. Weiss. 2018. ZAP-70 in Signaling, Biology, and Disease. Annual Review of Immunology. 36(1): p. 127–156.

12. Lin, J. and A. Weiss. 2001. Identification of the minimal tyrosine residues required for linker for activation of T cell function. J. Biol. Chem. 276(31): p. 29588–29595.

13. Zhang, W., R.P. Trible, M. Zhu, S.K. Liu, C.J. McGlade, and L.E. Samelson. 2000. Association of Grb2, Gads and Phospholipase C-γ1 with phosphorylated LAT tyrosine residues. J. Biol. Chem. 275(30): p. 23355–23361.

14. Zhu, M., E. Janssen, and W. Zhang. 2003. Minimal requirement of tyrosine residues of linker for activation of T cells in TCR signaling and thymocyte development. J Immunol. 170(1): p. 325–33.

15. Paz, P.E., S. Wang, H. Clarke, X. Lu, D. Stokoe, and A. Abo. 2001. Mapping the Zap-70 phosphorylation sites on LAT (linker for activation of T cells) required for recruitment and activation of signalling proteins in T cells. Biochem J. 356(Pt 2): p. 461–71.

16. Wardenburg, J.B., C. Fu, J.K. Jackman, H. Flotow, S.E. Wilkinson, D.H. Williams, R. Johnson, G. Kong, A.C. Chan, and P.R. Findell. 1996. Phosphorylation of SLP-76 by the ZAP-70 protein-tyrosine kinase is required for T-cell receptor function. J. Biol. Chem. 271(33): p. 19641–19644.

17. Fang, N., D.G. Motto, S.E. Ross, and G.A. Koretzky. 1996. Tyrosines 113, 128, and 145 of SLP-76 are required for optimal augmentation of NFAT promoter activity. J. Immunol. 157: p. 3769–3773.

18. Koretzky, G.A., F. Abtahian, and M.A. Silverman. 2006. SLP76 and SLP65: complex regulation of signalling in lymphocytes and beyond. Nat. Rev. Immunol. 6: p. 67–78.

19. Heyeck, S.D., H.M. Wilcox, S.C. Bunnell, and L.J. Berg. 1997. Lck phosphorylates the activation loop tyrosine of the Itk kinase domain and activates Itk kinase activity. J Biol Chem. 272(40): p. 25401–8.

20. Bogin, Y., C. Ainey, D. Beach, and D. Yablonski. 2007. SLP-76 mediates and maintains activation of the Tec family kinase ITK via the T cell antigen receptor-induced association between SLP-76 and ITK. Proc. Natl. Acad. Sci. USA. 104(16): p. 6638–43.

21. Sela, M., Y. Bogin, D. Beach, T. Oellerich, J. Lehne, J.E. Smith-Garvin, M. Okumura, E. Starosvetsky, R. Kosoff, E. Libman, G. Koretzky, T. Kambayashi, H. Urlaub, J. Wienands, J. Chernoff, and D. Yablonski. 2011. Sequential phosphorylation of SLP-76 at tyrosine 173 is required for activation of T and mast cells. EMBO J. 30(15): p. 3160–72.

22. Devkota, S., R.E. Joseph, L. Min, D. Bruce Fulton, and A.H. Andreotti. 2015. Scaffold Protein SLP-76 Primes PLCγ1 for Activation by ITK-Mediated Phosphorylation. Journal of Molecular Biology. 427(17): p. 2734–47.

23. Poulin, B., F. Sekiya, and S.G. Rhee. 2005. Intramolecular interaction between phosphorylated tyrosine-783 and the C-terminal Src homology 2 domain activates phospholipase C-gamma1. Proc. Natl. Acad. Sci. USA. 102(12): p. 4276–81.

24. Andreotti, A.H., S.C. Bunnell, S. Feng, L.J. Berg, and S.L. Schreiber. 1997. Regulatory intramolecular association in a tyrosine kinase of the Tec family. Nature. 385(6611): p. 93–7.

25. Andreotti, A.H., P.L. Schwartzberg, R.E. Joseph, and L.J. Berg. 2010. T-Cell Signaling Regulated by the Tec Family Kinase, Itk. Cold Spring Harbor Perspectives in Biology. 2(7): p. a002287.

26. Devkota, S., R.E. Joseph, S.E. Boyken, D.B. Fulton, and A.H. Andreotti. 2017. An Autoinhibitory Role for the Pleckstrin Homology Domain of Interleukin-2-Inducible Tyrosine Kinase and Its Interplay with Canonical Phospholipid Recognition. Biochemistry. 56(23): p. 2938–2949.

27. Su, Y.-W., Y. Zhang, J. Schweikert, G.A. Koretzky, M. Reth, and J. Wienands. 1999. Interaction of SLP adaptors with the SH2 domain of Tec family kinases. Eur. J. Immunol. 29: p. 3702–3711.

28. Bunnell, S.C., M. Diehn, M.B. Yaffe, P.R. Findell, L.C. Cantley, and L.J. Berg. 2000. Biochemical interactions integrating Itk with the T Cell Receptor-initiated signaling cascade. J. Biol. Chem. 275(3): p. 2219–2230.

29. Grasis, J.A., D.M. Guimond, N.R. Cam, K. Herman, P. Magotti, J.D. Lambris, and C.D. Tsoukas. 2010. In Vivo Significance of ITK-SLP-76 Interaction in Cytokine Production. Mol. Cell. Biol. 30(14): p. 3596–3609.

30. Jordan, M.S., J.E. Smith, J.C. Burns, J.E. Austin, K.E. Nichols, A.C. Aschenbrenner, and G.A. Koretzky. 2008. Complementation in trans of altered thymocyte development in mice expressing mutant forms of the adaptor molecule SLP76. Immunity. 28(3): p. 359–69.

31. Jordan, M.S. and G.A. Koretzky. 2010. Coordination of receptor signaling in multiple hematopoietic cell lineages by the adaptor protein SLP-76. Cold Spring Harb Perspect Biol. 2(4): p. a002501.

32. Gonen, R., D. Beach, C. Ainey, and D. Yablonski. 2005. T Cell Receptor-induced Activation of Phospholipase C-γ1 Depends on a Sequence-independent Function of the P-I Region of SLP-76. J. Biol. Chem. 280(9): p. 8364–70.

33. Houtman, J.C., Y. Higashimoto, N. Dimasi, S. Cho, H. Yamaguchi, B. Bowden, C. Regan, E.L. Malchiodi, R. Mariuzza, P. Schuck, E. Appella, and L.E. Samelson. 2004. Binding specificity of multiprotein signaling complexes is determined by both cooperative interactions and affinity preferences. Biochemistry. 43(14): p. 4170–8.

34. Kumar, L., S. Feske, A. Rao, and R.S. Geha. 2005. A 10-aa-long sequence in SLP-76 upstream of the Gads binding site is essential for T cell development and function. Proc Natl Acad Sci U S A. 102(52): p. 19063–8.

35. Sanzenbacher, R., D. Kabelitz, and O. Janssen. 1999. SLP-76 Binding to p56^lck^: A role for SLP-76 in CD4-induced desensitization of the TCR/CD3 signaling complex. J. Immunol. 163: p. 3143–3152.

36. Liu, K.-Q., S.C. Bunnell, C.B. Gurniak, and L.J. Berg. 1998. T cell receptor-initiated calcium release is uncoupled from capacitative calcium entry in Itk-deficient T cells. J. Exp. Med. 187(10): p. 1721–1727.

37. Fowell, D.J., K. Shinkai, X.C. Liao, A.M. Beebe, R.L. Coffman, D.R. Littman, and R.M. Locksley. 1999. Impaired NFATc translocation and failure of Th2 development in Itk-deficient CD4+ T cells. Immunity. 11(4): p. 399–409.

38. Lugassy, J., J. Corso, D. Beach, T. Petrik, T. Oellerich, H. Urlaub, and D. Yablonski. 2015. Modulation of TCR responsiveness by the Grb2-family adaptor, Gads. Cell Signal. 27(1): p. 125–134.

39. Bilal, M.Y., E.Y. Zhang, B. Dinkel, D. Hardy, T.M. Yankee, and J.C. Houtman. 2015. GADS is required for TCR-mediated calcium influx and cytokine release, but not cellular adhesion, in human T cells. Cell Signal. 27(4): p. 841–50.

40. Berry, D.M., P. Nash, S.K. Liu, T. Pawson, and C.J. McGlade. 2002. A high-affinity Arg-X-X-Lys SH3 binding motif confers specificity for the interaction between Gads and SLP-76 in T cell signaling. Curr Biol. 12(15): p. 1336–41.

41. Seet, B.T., D.M. Berry, J.S. Maltzman, J. Shabason, M. Raina, G.A. Koretzky, C.J. McGlade, and T. Pawson. 2007. Efficient T-cell receptor signaling requires a high-affinity interaction between the Gads C-SH3 domain and the SLP-76 RxxK motif. Embo J. 26(3): p. 678–689.

42. Sukenik, S., M.P. Frushicheva, C. Waknin-Lellouche, E. Hallumi, T. Ifrach, R. Shalah, D. Beach, R. Avidan, I. Oz, E. Libman, A. Aronheim, O. Lewinson, and D. Yablonski. 2017. Dimerization of the adaptor Gads facilitates antigen receptor signaling by promoting the cooperative binding of Gads to the adaptor LAT. Sci Signal. 10(498): p. eaal1482.

43. Di Bartolo, V., B. Montagne, M. Salek, B. Jungwirth, F. Carrette, J. Fourtane, N. Sol-Foulon, F. Michel, O. Schwartz, W.D. Lehmann, and O. Acuto. 2007. A novel pathway down-modulating T cell activation involves HPK-1-dependent recruitment of 14-3-3 proteins on SLP-76. J. Exp. Med. 204(3): p. 681–91.

44. Shui, J.W., J.S. Boomer, J. Han, J. Xu, G.A. Dement, G. Zhou, and T.H. Tan. 2007. Hematopoietic progenitor kinase 1 negatively regulates T cell receptor signaling and T cell-mediated immune responses. Nat. Immunol. 8(1): p. 84–91.

45. Lasserre, R., C. Cuche, R. Blecher-Gonen, E. Libman, E. Biquand, A. Danckaert, D. Yablonski, A. Alcover, and V. Di Bartolo. 2011. Release of serine/threonine-phosphorylated adaptors from signaling microclusters downregulates T cell activation. J. Cell. Biol. 195(5): p. 839–53.

46. Joseph, R.E., L. Min, R. Xu, E.D. Musselman, and A.H. Andreotti. 2007. A remote substrate docking mechanism for the tec family tyrosine kinases. Biochemistry. 46(18): p. 5595–603.

47. Min, L., R.E. Joseph, D.B. Fulton, and A.H. Andreotti. 2009. Itk tyrosine kinase substrate docking is mediated by a nonclassical SH2 domain surface of PLCgamma1. Proc. Natl. Acad. Sci. USA. 106(50): p. 21143–8.

48. Tomlinson, M.G., T. Kurosaki, A.E. Berson, G.H. Fujii, J.A. Johnston, and J.B. Bolen. 1999. Reconstitution of Btk signaling by the atypical tec family tyrosine kinases Bmx and Txk. J Biol Chem. 274(19): p. 13577–85.

49. Weiss, A. and J.D. Stobo. 1984. Requirement for the coexpression of T3 and the T cell antigen receptor on a malignant human T cell line. J. Exp. Med. 160: p. 1284–1299.

50. Schmidt, T.G.M., L. Batz, L. Bonet, U. Carl, G. Holzapfel, K. Kiem, K. Matulewicz, D. Niermeier, I. Schuchardt, and K. Stanar. 2013. Development of the Twin-Strep-tag® and its application for purification of recombinant proteins from cell culture supernatants. Protein Expression and Purification. 92(1): p. 54–61.

51. Pear, W.S., J.P. Miller, L. Xu, J.C. Pui, B. Soffer, R.C. Quackenbush, A.M. Pendergast, R. Bronson, J.C. Aster, M.L. Scott, and D. Baltimore. 1998. Efficient and rapid induction of a chronic myelogenous leukemia-like myeloproliferative disease in mice receiving P210 bcr/abl-transduced bone marrow. Blood. 92(10): p. 3780–92.

52. Day, R.N. and M.W. Davidson. 2009. The fluorescent protein palette: tools for cellular imaging. Chemical Society Reviews. 38(10): p. 2887–2921.

53. Bourgin, C., R.P. Bourette, S. Arnaud, Y. Liu, L.R. Rohrschneider, and G. Mouchiroud. 2002. Induced Expression and Association of the Mona/Gads Adapter and Gab3 Scaffolding Protein during Monocyte/Macrophage Differentiation. Molecular and Cellular Biology. 22(11): p. 3744–3756.

54. Yablonski, D., M.R. Kuhne, T. Kadlecek, and A. Weiss. 1998. Uncoupling of nonreceptor tyrosine kinases from PLC-γ1 in an SLP-76-deficient T cell. Science. 281: p. 413–416.

55. Lo, W.-L., N.H. Shah, S.A. Rubin, W. Zhang, V. Horkova, I.R. Fallahee, O. Stepanek, L.I. Zon, J. Kuriyan, and A. Weiss. 2019. Slow phosphorylation of a tyrosine residue in LAT optimizes T cell ligand discrimination. Nature Immunology. 20(11): p. 1481–1493.

56. Lo, W.-L., N.H. Shah, N. Ahsan, V. Horkova, O. Stepanek, A.R. Salomon, J. Kuriyan, and A. Weiss. 2018. Lck promotes Zap70-dependent LAT phosphorylation by bridging Zap70 to LAT. Nature Immunology. 19(7): p. 733–741.

57. Oellerich, T., M. Gronborg, K. Neumann, H.H. Hsiao, H. Urlaub, and J. Wienands. 2009. SLP-65 phosphorylation dynamics reveals a functional basis for signal integration by receptor-proximal adaptor proteins. Mol. Cell. Proteomics. 8(7): p. 1738–50.

58. Hornbeck, P.V., B. Zhang, B. Murray, J.M. Kornhauser, V. Latham, and E. Skrzypek. 2014. PhosphoSitePlus, 2014: mutations, PTMs and recalibrations. Nucleic Acids Research. 43(D1): p. D512–D520.

59. Mayya, V., D.H. Lundgren, S.I. Hwang, K. Rezaul, L. Wu, J.K. Eng, V. Rodionov, and D.K. Han. 2009. Quantitative phosphoproteomic analysis of T cell receptor signaling reveals system-wide modulation of protein-protein interactions. Sci Signal. 2(84): p. ra46.

60. Kim, J.-E. and F.M. White. 2006. Quantitative Analysis of Phosphotyrosine Signaling Networks Triggered by CD3 and CD28 Costimulation in Jurkat Cells. The Journal of Immunology. 176(5): p. 2833–2843.

61. Ross, Sarah H., C. Rollings, Karen E. Anderson, Phillip T. Hawkins, Len R. Stephens, and Doreen A. Cantrell. 2016. Phosphoproteomic Analyses of Interleukin 2 Signaling Reveal Integrated JAK Kinase-Dependent and - Independent Networks in CD8^+^ T Cells. Immunity. 45(3): p. 685–700.

62. Sievers, F., A. Wilm, D. Dineen, T.J. Gibson, K. Karplus, W. Li, R. Lopez, H. McWilliam, M. Remmert, J. Soding, J.D. Thompson, and D.G. Higgins. 2011. Fast, scalable generation of high-quality protein multiple sequence alignments using Clustal Omega. Mol Syst Biol. 7: p. 539.

63. Crooks, G.E., G. Hon, J.M. Chandonia, and S.E. Brenner. 2004. WebLogo: a sequence logo generator. Genome Res. 14(6): p. 1188–90.

64. Cruz-Orcutt, N., A. Vacaflores, S.F. Connolly, S.C. Bunnell, and J.C.D. Houtman. 2014. Activated PLC-γ1 is catalytically induced at LAT but activated PLC-γ1 is localized at both LAT- and TCR-containing complexes. Cell Signal. 26(4): p. 797–805.

65. Shah, N.H., Q. Wang, Q. Yan, D. Karandur, T.A. Kadlecek, I.R. Fallahee, W.P. Russ, R. Ranganathan, A. Weiss, and J. Kuriyan. 2016. An electrostatic selection mechanism controls sequential kinase signaling downstream of the T cell receptor. eLife. 5: p. e20105.

66. Wilcox, H.M. and L.J. Berg. 2003. Itk phosphorylation sites are required for functional activity in primary T cells. J Biol Chem. 278(39): p. 37112–21.

67. van de Weyer, P.S., M. Muehlfeit, C. Klose, J.V. Bonventre, G. Walz, and E.W. Kuehn. 2006. A highly conserved tyrosine of Tim-3 is phosphorylated upon stimulation by its ligand galectin-9. Biochemical and Biophysical Research Communications. 351(2): p. 571–576.

68. Hwang, E.S., S.J. Szabo, P.L. Schwartzberg, and L.H. Glimcher. 2005. T Helper Cell Fate Specified by Kinase-Mediated Interaction of T-bet with GATA-3. Science. 307(5708): p. 430–433.

69. Hey, F., N. Czyzewicz, P. Jones, and F. Sablitzky. 2012. DEF6, a Novel Substrate for the Tec Kinase ITK, Contains a Glutamine-rich Aggregation-prone Region and Forms Cytoplasmic Granules that Co-localize with P-bodies. Journal of Biological Chemistry. 287(37): p. 31073–31084.

70. Lin, T.A., K.W. McIntyre, J. Das, C. Liu, K.D. O’Day, B. Penhallow, C.Y. Hung, G.S. Whitney, D.J. Shuster, X. Yang, R. Townsend, J. Postelnek, S.H. Spergel, J. Lin, R.V. Moquin, J.A. Furch, A.V. Kamath, H. Zhang, P.H. Marathe, J.J. Perez-Villar, A. Doweyko, L. Killar, J.H. Dodd, J.C. Barrish, J. Wityak, and S.B. Kanner. 2004. Selective Itk inhibitors block T-cell activation and murine lung inflammation. Biochemistry. 43(34): p. 11056–62.

71. Liu, S.K., N. Fang, G.A. Koretzky, and C.J. McGlade. 1999. The hematopoietic-specific adaptor protein Gads functions in T-cell signaling via interactions with the SLP-76 and LAT adaptors. Curr. Biol. 9: p. 67–75.

72. Asada, H., N. Ishii, Y. Sasaki, K. Endo, H. Kasai, N. Tanaka, T. Takeshita, S. Tsuchiya, T. Konno, and K. Sugamura. 1999. Grf40, a novel Grb2 family member, is involved in T cell signaling through interactions with SLP-76 and LAT. J. Exp. Med. 189(9): p. 1383–1390.

73. Law, C.-L., M.K. Ewings, P.M. Chaudhary, S.A. Solow, T.J. Yun, A.J. Marshall, L. Hood, and E.A. Clark. 1999. GrpL, a Grb2-related adaptor protein, interacts with SLP-76 to regulate Nuclear Factor of Activated T Cell activation. J. Exp. Med. 189(8): p. 1243–1253.

74. Brazin, K.N., D.B. Fulton, and A.H. Andreotti. 2000. A specific intermolecular association between the regulatory domains of a Tec family kinase. J Mol Biol. 302(3): p. 607–23.

75. Dombroski, D., R.A. Houghtling, C.M. Labno, P. Precht, A. Takesono, N.J. Caplen, D.D. Billadeau, R.L. Wange, J.K. Burkhardt, and P.L. Schwartzberg. 2005. Kinase-independent functions for Itk in TCR-induced regulation of Vav and the actin cytoskeleton. J. Immunol. 174(3): p. 1385–92.

76. Yoder, J., C. Pham, Y.-M. Iizuka, O. Kanagawa, S.K. Liu, J. McGlade, and A.M. Cheng. 2001. Requirement for the SLP-76 adaptor GADS in T cell development. Science. 291: p. 1987–1991.

77. Tatárová, Z., J. Brábek, D. Rösel, and M. Novotný. 2012. SH3 Domain Tyrosine Phosphorylation – Sites, Role and Evolution. PLoS ONE. 7(5): p. e36310.

78. Merő, B., L. Radnai, G. Gógl, O. Tőke, I. Leveles, K. Koprivanacz, B. Szeder, M. Dülk, G. Kudlik, V. Vas, A. Cserkaszky, S. Sipeki, L. Nyitray, B.G. Vértessy, and L. Buday. 2019. Structural insights into the tyrosine phosphorylation–mediated inhibition of SH3 domain–ligand interactions. Journal of Biological Chemistry. 294(12): p. 4608–4620.

79. Broome, M.A. and T. Hunter. 1996. Requirement for c-Src catalytic activity and the SH3 domain in platelet-derived growth factor BB and epidermal growth factor mitogenic signaling. J Biol Chem. 271(28): p. 16798–806.

80. Pesti, S., A. Balazs, R. Udupa, B. Szabo, A. Fekete, G. Bogel, and L. Buday. 2012. Complex formation of EphB1/Nck/Caskin1 leads to tyrosine phosphorylation and structural changes of the Caskin1 SH3 domain. Cell Commun Signal. 10(1): p. 36.

81. Andersen, T.C.B., P.E. Kristiansen, Z. Huszenicza, M.U. Johansson, R.P. Gopalakrishnan, H. Kjelstrup, S. Boyken, V. Sundvold-Gjerstad, S. Granum, M. Sorli, P.H. Backe, D.B. Fulton, B.G. Karlsson, A.H. Andreotti, and A. Spurkland. 2019. The SH3 domains of the protein kinases ITK and LCK compete for adjacent sites on T cell-specific adapter protein. J Biol Chem. 294(42): p. 15480–15494.

82. Joseph, R.E., D.B. Fulton, and A.H. Andreotti. 2007. Mechanism and functional significance of Itk autophosphorylation. J Mol Biol. 373(5): p. 1281–92.

83. Irvin, B.J., B. Williams, L., A.E. Nilson, H.O. Maynor, and R.T. Abraham. 2000. Pleiotropic contributions of phospholipase C-γ1 (PLC-γ1) to T-cell antigen receptor-mediated signaling: reconstitution studies of a PLC-γ1-deficient Jurkat T-cell line. Mol. Cell. Biol. 20(24): p. 9149–9161.

84. Sauer, K., J. Liou, S.B. Singh, D. Yablonski, A. Weiss, and R.M. Perlmutter. 2001. Hematopoietic Progenitor Kinase 1 associates physically and functionally with the adaptor proteins B Cell Linker Protein and SLP-76 in lymphocytes. J. Biol. Chem. 276: p. 45207–45216.

85. Brenner, D., M. Brechmann, S. Rohling, M. Tapernoux, T. Mock, D. Winter, W.D. Lehmann, F. Kiefer, M. Thome, P.H. Krammer, and R. Arnold. 2009. Phosphorylation of CARMA1 by HPK1 is critical for NF-kappaB activation in T cells. Proc Natl Acad Sci U S A. 106(34): p. 14508–13.

86. Liu, H., H. Schneider, A. Recino, C. Richardson, M.W. Goldberg, and C.E. Rudd. 2015. The Immune Adaptor SLP-76 Binds to SUMO-RANGAP1 at Nuclear Pore Complex Filaments to Regulate Nuclear Import of Transcription Factors in T Cells. Mol Cell. 59(5): p. 840–9.

87. Ellis, J.H., C. Ashman, M.N. Burden, K.E. Kilpatrick, M.A. Morse, and P.A. Hamblin. 2000. GRID: a novel Grb-2-related adapter protein that interacts with the activated T cell costimulatory receptor CD28. J. Immunol. 164: p. 5805–5814.

88. Watanabe, R., Y. Harada, K. Takeda, J. Takahashi, K. Ohnuki, S. Ogawa, D. Ohgai, N. Kaibara, O. Koiwai, K. Tanabe, H. Toma, K. Sugamura, and R. Abe. 2006. Grb2 and Gads exhibit different interactions with CD28 and play distinct roles in CD28-mediated costimulation. J Immunol. 177(2): p. 1085–91.

89. Higo, K., M. Oda, H. Morii, J. Takahashi, Y. Harada, S. Ogawa, and R. Abe. 2014. Quantitative analysis by surface plasmon resonance of CD28 interaction with cytoplasmic adaptor molecules Grb2, Gads and p85 PI3K. Immunological Investigations. 43(3): p. 278–291.

90. Thaker, Y.R., H. Schneider, and C.E. Rudd. 2015. TCR and CD28 activate the transcription factor NF-κB in T-cells via distinct adaptor signaling complexes. Immunology Letters. 163(1): p. 113–119.

91. Greene, T.A., P. Powell, C. Nzerem, M.J. Shapiro, and V.S. Shapiro. 2003. Cloning and characterization of ALX, an adaptor downstream of CD28. J Biol Chem. 278(46): p. 45128–34.

92. Shapiro, M.J., P. Powell, A. Ndubuizu, C. Nzerem, and V.S. Shapiro. 2004. The ALX Src Homology 2 Domain Is Both Necessary and Sufficient to Inhibit T Cell receptor/CD28-mediated Up-regulation of RE/AP. Journal of Biological Chemistry. 279(39): p. 40647–40652.

