## Supplementary Figures for "Itk promotes the integration of TCR and CD28 costimulation, through its direct substrates, SLP-76 and Gads"

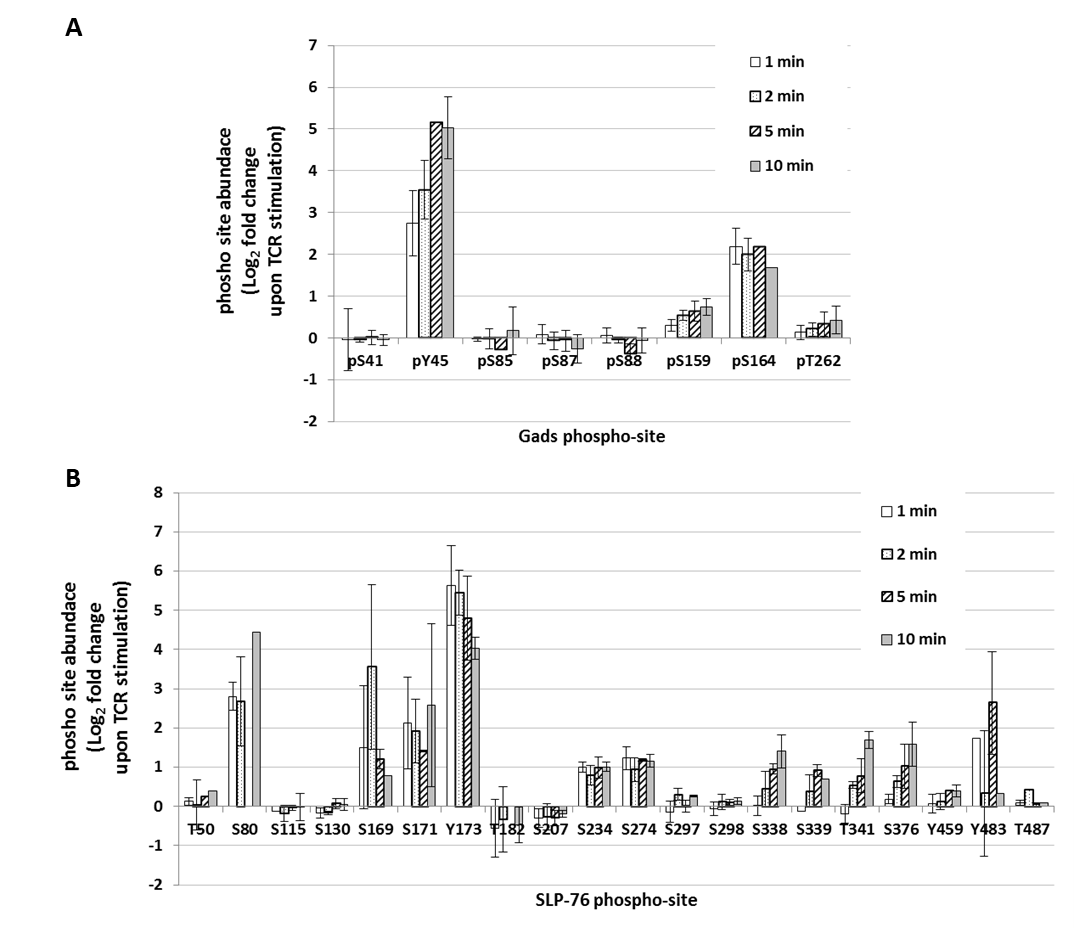

**Figure S1. Summary of SLP-76 and Gads phosphorylation sites identified in our SILAC-based phospho-mass spectrometry analysis.** A SILAC-based approach was used to identify the TCR-induced change in site-specific phosphorylation of Gads (**A**) and SLP-76 (**B**), as described in Materials and Methods. Four biological replicates were performed. Here, we present the median Log_2_-fold change in abundance, observed upon TCR stimulation, for all Class I sites that were identified in at least two time points of at least two biological replicates. Certain well-established sites, such as SLP-76 Y113, Y128 and Y145 were not identified in our experiments, due to the lack of a nearby trypsin, chymotrypsin or AspN site. Error bars indicate the SD.

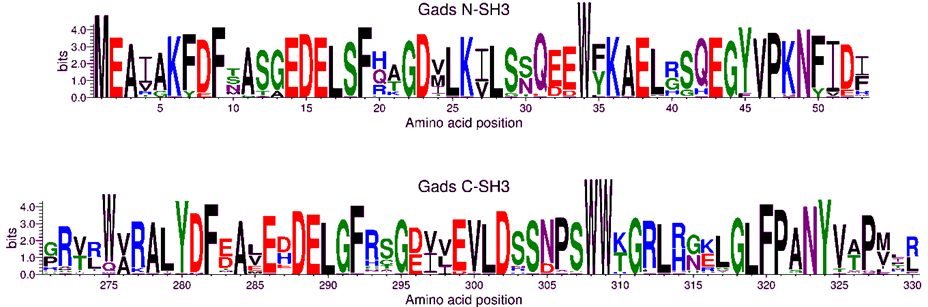

**Figure S2. Evolutionary conservation of the Gads N- and C-terminal SH3 domains.** 66 vertebrate Gads orthologs representing 55 different taxonomical orders were aligned, as described in Figure 2B, and WebLogo was used to depict sequence conservation within the N- and C-terminal SH3 domains. Amino acid numbering is according to the sequence of human Gads.

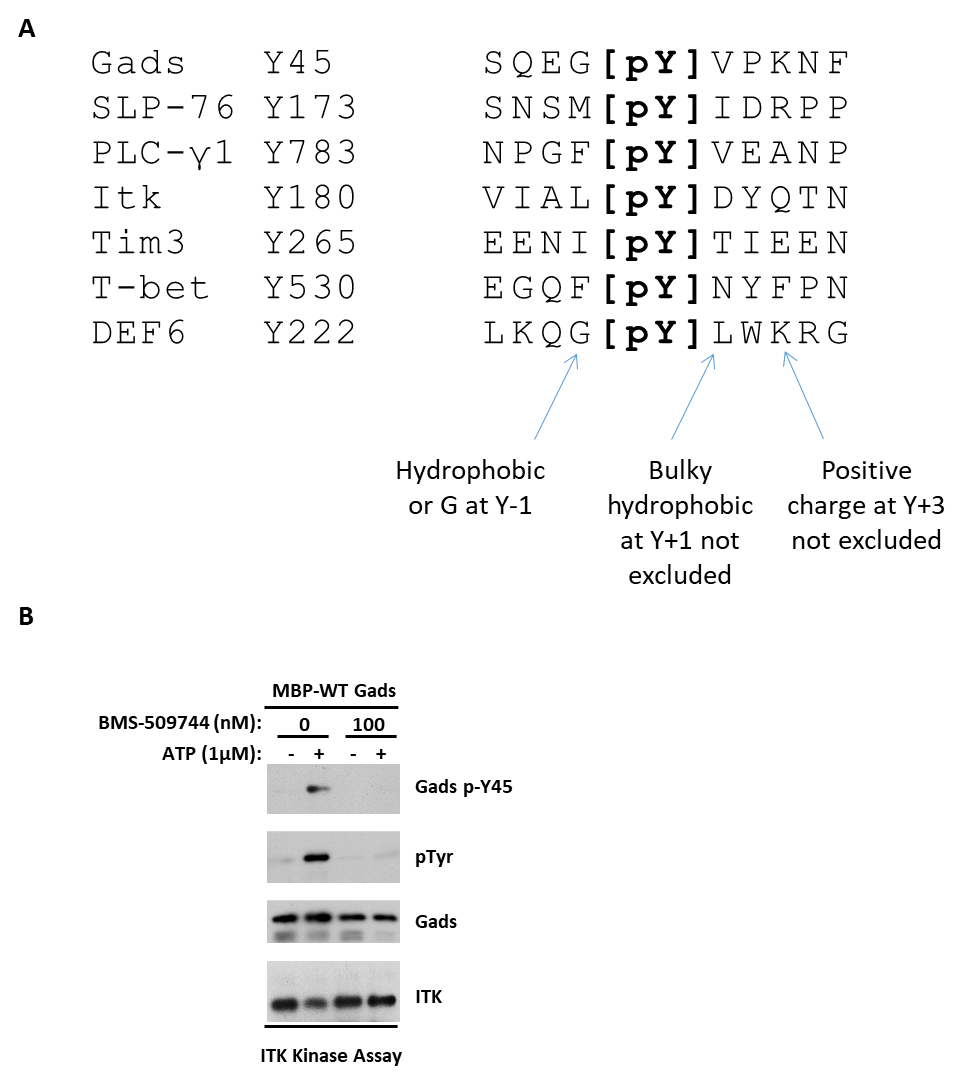

**Figure S3.** **Gads Y45 is an Itk-targeted site** (**A**) **Comparison of the Y45 motif to known ITK-targeted sites.** Alignment of Gads Y45 to previously-described Itk-mediated phosphorylation sites, including SLP-76 p-Y173, PLCγ1 p-Y783, Itk p-Y180, Tim3 p-Y265, T-bet p-Y530 (525 in mouse), and DEF6 p-Y222. Some shared characteristics are noted. (**B**) **Specificity of bead-bound Itk preparation.** *in vitro* phosphorylation of Gads by Itk was performed as in Figure 4A, using WT Gads as a substrate, in the presence or absence of the ITK-specific inhibitor, BMS-509744.

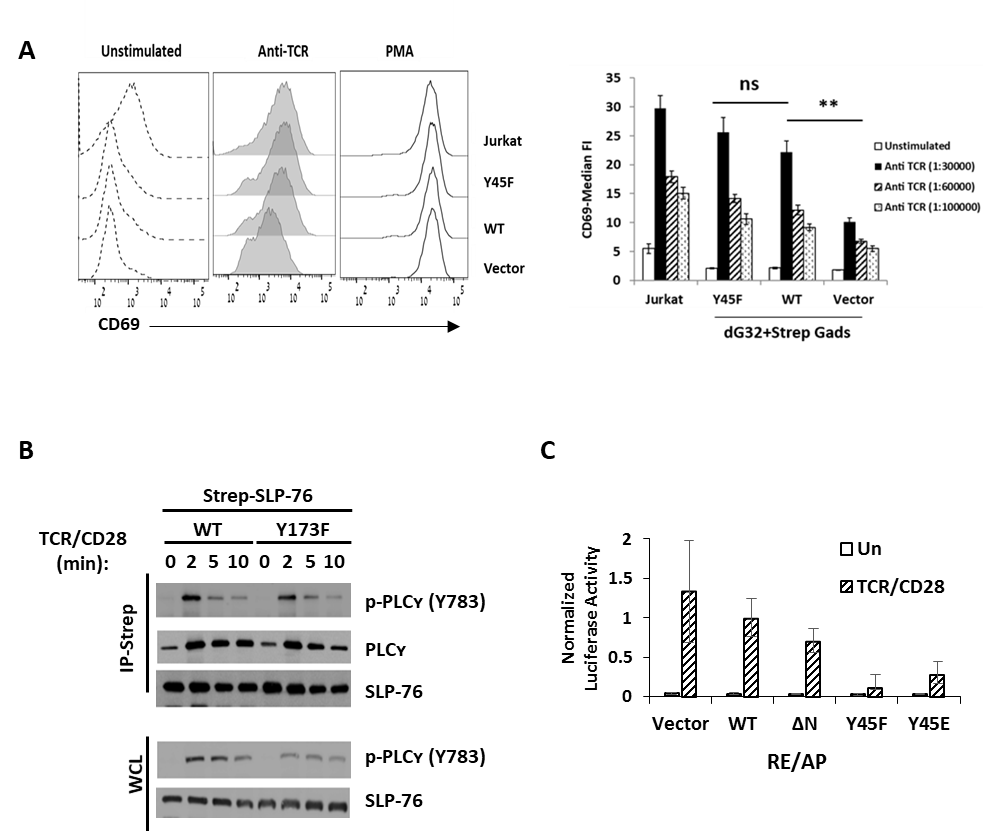

**Figure S4. Downstream signaling functions of Gads p-Y45 and SLP-76 p-173.** (**A**) **CD69 response is independent of Gads Y45.** A FACS-based assay was used to measure TCR-induced CD69 expression in Jurkat, dG32 or dG32 cells that were stably reconstituted with the indicated twin strep-tagged Gads alleles, expressed from an IRES-GFP-marked retroviral vector. Cell lines were differentially barcoded with CellTrace Violet and mixed together, prior to stimulation. Cells were mock stimulated, or stimulated in triplicate overnight with the indicated concentrations of anti-TCR (C305) or phorbol 12-myristate 13-acetate (PMA), and then stained with anti-CD69. Results were analyzed while gating on matched GFP-expression gates. Left: Representative results when anti-TCR was used at 1:30,000. Right: Median TCR-induced CD69 cell surface abundance for each cell line was normalized to that induced by PMA (n=3; error bars indicate the SD). Statistical analysis was performed using the unpaired two-tailed *t*-test, comparing the TCR response of each of the indicated cell lines to the response of WT-reconstituted dG32 cells, with the same stimulus (**, p<0.005). (**B**) **SLP-76 pY173 mediates the release of phospho-PLC-γ1 from the LAT-nucleated complex.** J14 cells, stably reconstituted with twin strep-tagged SLP-76, either wild type or bearing the Y173F mutation were stimulated and lysed as in Figure 5B, and whole cell lysates (WCL), or streptactin-purified SLP-76 complexes were probed with the indicated phospho-specific or total protein antibodies. (**C**) **TCR/CD28-induced activation of RE/AP depends on Gads Y45.** dG32 cells were independently reconstituted with the indicated forms of twin-strep-tagged Gads, and were sorted for comparable expression level, thus creating a set of cell lines distinct from those used for Figure 8F. RE/AP luciferase activity was measured as in Figure 8F. Results are the average of 2 experiments, conducted in triplicate, error bars indicate the SD.
